# CPT1A loss promotes lung metastasis in immune-competent mice via a mechanism of mtDNA release and chronic activation of STING pathway

**DOI:** 10.64898/2026.05.01.722261

**Authors:** Xiaoyong Wang, Shu-Ting Chou, Yoonha Hwang, Jin Chen, Deanna N. Edwards

## Abstract

The metastatic progression of breast cancer involves complex interactions between tumor cells and immune cells, including T cells that exert cytotoxic pressure to limit metastasis. Tumor cells reprogram their metabolism to evade immune surveillance, a critical step to achieving metastatic outgrowth. Using an unbiased CRISPR screen targeting metabolism-related genes and a clinically relevant spontaneous metastasis mouse model, we identified CPT1A, the rate-limiting enzyme in fatty acid β-oxidation, as a suppressor of immune-dependent metastasis. Loss of CPT1A enhances lung metastasis in immunocompetent mice, but not Rag1 KO mice that lack mature lymphocytes. Loss of CPT1A triggers cytosolic mitochondrial DNA (mtDNA) release via the mPTP pore. Cytosolic mtDNA release triggers a STING-dependent inflammatory response, creating an environment that impairs CD8+ T cell function, promoting metastatic outgrowth. Among breast cancer patients, low *CPT1A* expression correlates with poor survival when CD8+ T cell infiltration is high. These findings reveal an extrinsic role for CPT1A in immune-tumor dynamics and suggest therapeutic opportunities targeting inflammation in metastatic breast cancer.

## Introduction

The metastatic cascade is a complex multifaceted process resulting in tumor cells disseminating to distant niches. During metastatic progression, tumor cells encounter immune cells and evasion of immunosurveillance mechanisms are necessary to complete metastatic outgrowth^1–5^. T cells play a decisive role in metastasis by exerting cytotoxic pressure on disseminated tumor cells^1–3^. Tumor cells reprogram their metabolism to modulate anti-tumor responses^6–9^, but the specific metabolic drivers that influence the crosstalk between metastasizing tumor cells and T cells remain poorly understood.

Fatty acid β-oxidation (FAO) is a critical metabolic source of ATP, NADPH, and acetyl-coA necessary to provide energy, protection from oxidation, and epigenetic regulation necessary for tumor growth and progression^10^. The rate limiting enzyme of FAO, carnitine palmitoyltransferase 1A (CPT1A), has been well characterized as a tumor cell intrinsic driver of tumor growth and metastatic progression^11–17^. However, inconsistent with this intrinsic role, highly aggressive breast cancer subtypes, including basal-like breast cancer, express lower levels of CPT1A than ER+ subtypes^11,18^. Indeed, CPT1A loss has been implicated in driving inflammation^19–22^, supporting a potential extrinsic role during metastasis. Given that immunosuppression builds gradually over time^23,24^, the extrinsic influence of CPT1A may be progressive in nature that is not effectively represented by most metastasis models. Therefore, the specific extrinsic role of CPT1A in breast cancer metastasis is unclear.

To evaluate extrinsic metabolic drivers of immune-dependent breast cancer metastasis, we utilized a gradual metastatic model that better recapitulates disease progression in patients, including surgical resection of the primary tumor followed by a latent period of metastatic progression. An in vivo loss of function screen identified CPT1A as a suppressor of metastatic progression. We show that loss of CPT1A in breast cancer cells enhances spontaneous metastasis to the lungs of immunocompetent, but not lymphocyte-deficient murine hosts. CPT1A deficiency drives a STING-dependent inflammatory response, instigated by cytosolic mitochondrial DNA (mtDNA) release through the mitochondrial permeability transition pore (mPTP) to create an immunosuppressive microenvironment at the metastatic site. In patients, low CPT1A expression predicts poor outcomes in immune-enriched populations. These findings describe a tumor cell extrinsic role of CPT1A on metastatic suppression in breast cancer, highlighting the complex relationship between metabolism and immunoregulation during metastasis.

## Results

### In vivo CRISPR screen reveals immune-dependent metabolic genes that regulate metastasis

We established a CRISPR screening approach to identify metabolic genes that alter the metastatic capacity of breast cancer cells. Cas9-expressing 4T1 were infected with a custom pooled CRISPR library of 404 sgRNA’s targeting 101 metabolism-related genes (Fig. 1a and Extended Data Fig. 1a). To evaluate immune-related contributors to metastasis, pooled cells were implanted into the inguinal mammary fat pad of female wild-type (WT) or Rag1^KO^ mice that lack B or T cells^25^ (Fig. 1a). We utilized a metastasis model that closely recapitulates the gradual progression of breast cancer in patients, including use of surgical resection followed by a period of metastatic outgrowth (Fig. 1a). Genomic DNA was collected from injection inputs, primary tumors at resection, and lung metastatic tumors (Fig. 1a). Library sgRNA representation was maintained across input and primary tumors, while fewer sgRNA’s were identified in lung metastatic tumors, consistent with the selective nature of metastasis (Fig. 1b and Extended Data Fig. 1b-d).

**Fig. 1.**
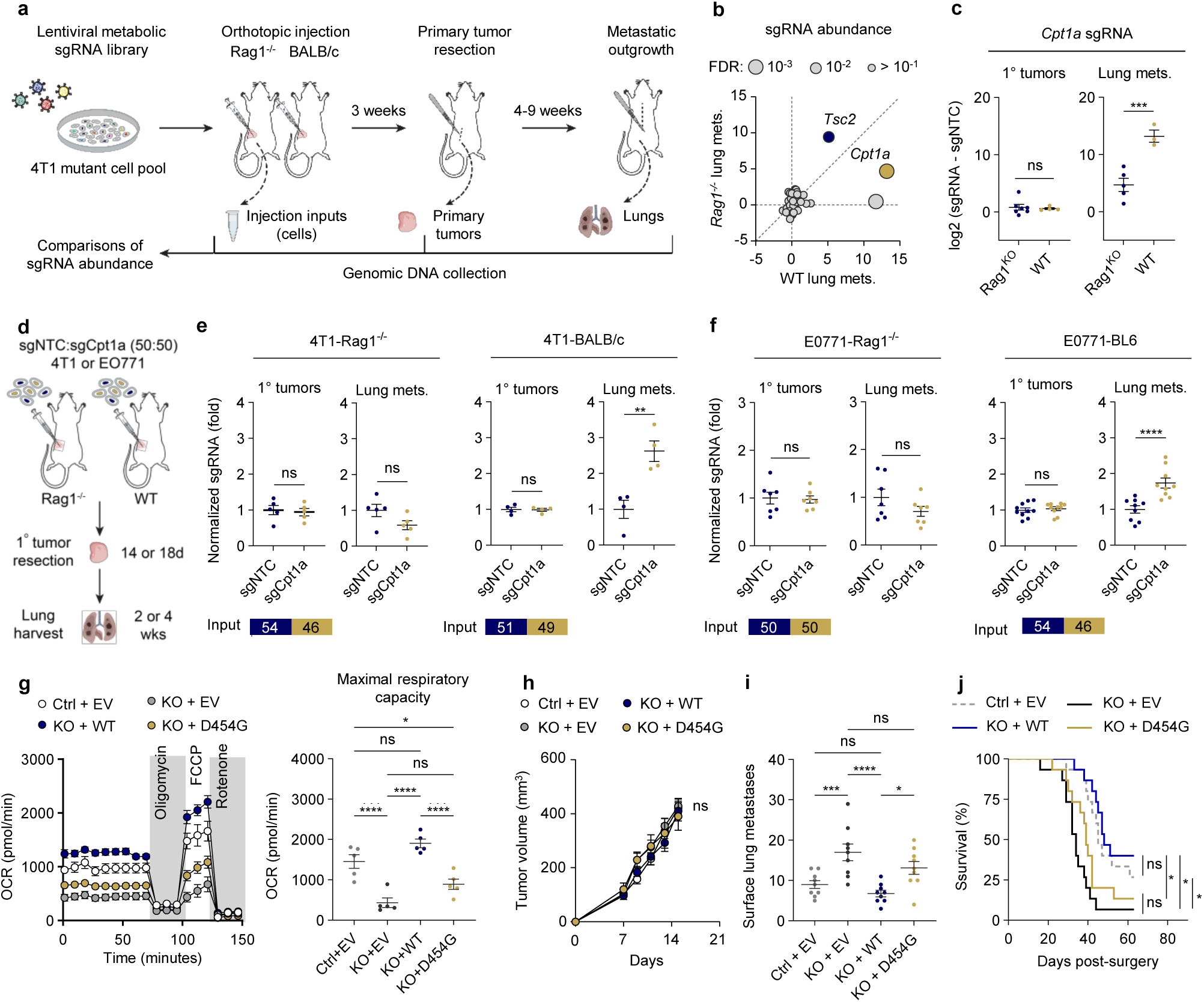
Cpt1a deficiency promotes breast cancer metastasis to lung. **a**, 4T1 cells transduced with metabolism-focused CRISPR KO lentiviral library were implanted into the mammary fat pad (MFP) of BALB/c Rag1^-/-^ or BALB/c wild type mice. Primary tumors were surgically removed on day 21 after transplantation and whole lung was collected (4-9 weeks). *n* ≥ 3 mice per group. **b**, The differential abundance (log_2_FC) of sgRNA corresponding genes in the lungs between *Rag1^-/-^* and wild type (WT) mice. Target gene sgRNAs were normalized to non-targeted control sgRNA (sgNTC). Cpt1a was enriched in WT mice. **c**, The log_2_-transformed Cpt1a sgRNA reads from the primary tumors (1° tumors) or lung metastases (lung mets.) between *Rag1*^-/-^ and WT mice. Welch’s t-test, p=0.688 (1° tumors), p=0.00199 (lung mets.) **d**, Diagram of *in vivo* metastasis competition assay in *Rag1*^-/-^ or BALB/c WT mice. Primary tumors were resected on day 14 (4T1) or day 18 (EO771) post-implantation, and whole lungs were collected 2 weeks and 4 weeks after 4T1, EO771 tumor resection, respectively. **e-f**, Normalized Cpt1a sgRNA (fold change) from primary tumors or lung metastases upon MFP injection of sgNTC control, or *Cpt1a*-KO 4T1 (**e**) or EO771 (**f**) into immune competent hosts (BALB/c) or immune deficient hosts (*Rag1^-/-^*). *n* ≥ 4 mice per group. Proportions of sgNTC vs s sgCpt1a cells at implantation are shown (Input). Welch’s t-test was performed. **g**, Oxygen consumption rate (pmol/min) in sgNTC control (Ctrl), *Cpt1a*-KO (KO) 4T1 cells or KO cells re-expressing WT or D454G CPT1A, per 5×10^4^ cells. Maximal respiration capacity OCR is shown. One-way ANOVA (p=4.32×10^-6^) with Tukey’s post hoc. **h-j,** 4T1 control, *Cpt1a*-KO, or re-expression WT or D454G CPT1A in *Cpt1*a-KO cells were implanted into the mammary fat pad of BALB/c mice (n=9), followed by primary tumor resection and lung metastatic growth. (**h**) Primary tumor volume was tracked over time. Two-way ANOVA (p=0.345). (**i**) Numbers of lung surface metastases at was determined at sacrifice. One-way ANOVA (p=1.17×10^-4^) with Tukey’s post-hoc. (**j**) Survival of mice inoculated with 4T1 control, *Cpt1a*-KO, or re-expression WT or D454G CPT1A in *Cpt1*a-KO tumors (n=15). *p<0.05, **p<0.01, ***p<0.005, ****p<0.001. ns, not significant.

Lung metastatic tumors from immunocompetent WT animals were highly correlative in sgRNA content, a stark contrast with primary tumors and Rag1^KO^ metastatic lesions (Extended Data Fig. 1b). The lack of variability was due to the significant enrichment of CPT1A-depeleted cells in WT but not Rag1^KO^ lung metastatic tumors (Fig. 1b-c and Extended Data Fig. 1e). Alternatively, Rag1^KO^ metastases were enriched with sgRNAs targeting the tumor suppressor Tsc2 (Fig. 1b and Extended Data Fig. 1f-g). Enrichment of sgCpt1a was consistent across all four sequences assessed in WT metastatic tumors, while no differences were observed in primary mammary tumors across models (Fig. 1c and Extended Data Fig. 1g), suggesting that loss of CPT1A increases metastasis in an immune-dependent manner.

### CPT1A enzymatic activity suppresses immune-dependent lung metastasis

To confirm that CPT1A loss promotes metastasis in the presence of an intact immune system, we generated Cpt1a knockout (KO) murine breast cancer cell lines (Extended Data Fig. 2a-b). Defective cellular respiration was observed in Cpt1a KO cells as well as reduced carnitine-loaded long-chain fatty acids, while other metabolic pathways were not affected (Extended Data Fig. 2c-g). To evaluate the metastatic capacity of Cpt1a KO cells, we utilized an unbiased, competitive spontaneous metastasis model that mimics tumor heterogeneity (Fig. 1d). NGS sequencing revealed that both 4T1 and E0771 sgCpt1a cells were more significantly enriched in lung metastatic tumors from immunocompetent mice than control cells, while lymphocyte deficiency eliminated or dramatically reduced this enrichment (Fig. 1e-f). However, control and Cpt1a KO cells were evenly distributed in the primary mammary tumors and cell viability was not significantly affected in multiple breast cancer cell lines (Fig. 1e-f and Extended Data Fig. 2h-i), indicating a specific role of CPT1A in metastatic progression to the lung.

Implantation of homogenous tumors into the mammary fat pad revealed an increase in spontaneous metastatic tumor burden of Cpt1a KO cells, although the primary tumor volume is unchanged from controls (Extended Data Fig. 2j-m). Thus, CPT1A offers protection against lung metastasis in an immune-dependent manner.

We next investigated whether the catalytic activity of CPT1A is responsible for metastatic suppression (Fig. 1g-j and Extended Data Fig. 2n-p). Several naturally occurring mutations have been described in CPT1A deficiency, a rare mitochondrial fatty acid oxidation disorder, including a D454G mutation within the carnitine acyl-transferase domain that results in loss of CPT1A enzymatic activity^26,27^ (Extended Data Fig. 2n). WT (Cpt1a^WT^) and D454G mutant (Cpt1a^D454G^) expression vectors were generated and expressed in Cpt1a KO breast cancer cells (Extended Data Fig. 2o).

Expression of Cpt1a^WT^ but not the enzyme-dead Cpt1a^D454G^ mutant rescued the fatty acid oxidation defect in Cpt1a KO cells, confirming the D454G mutation diminishes CPT1A catalytic activity (Fig. 1g and Extended Data Fig. 2p). In immunocompetent Cas9-tolerant mice, spontaneous lung metastasis was significantly reduced upon re-expression of Cpt1a^WT^, which was also associated with improved survival (Fig. 1h-j). However, the Cpt1a^D454G^ mutant did not rescue lung metastatic burden or survival (Fig. 1h-j), indicating the catalytic activity of CPT1A is necessary to suppress lung metastasis.

### CPT1A deficiency produces an inflammatory response through cytosolic mitochondrial DNA

To better understand how CPT1A loss may promote immune-dependent metastasis, we performed transcriptional analysis of control and Cpt1a KO cells. Inflammatory cytokines and chemokines were upregulated in Cpt1a KO breast cancer cells (Fig. 2).

**Fig. 2.**
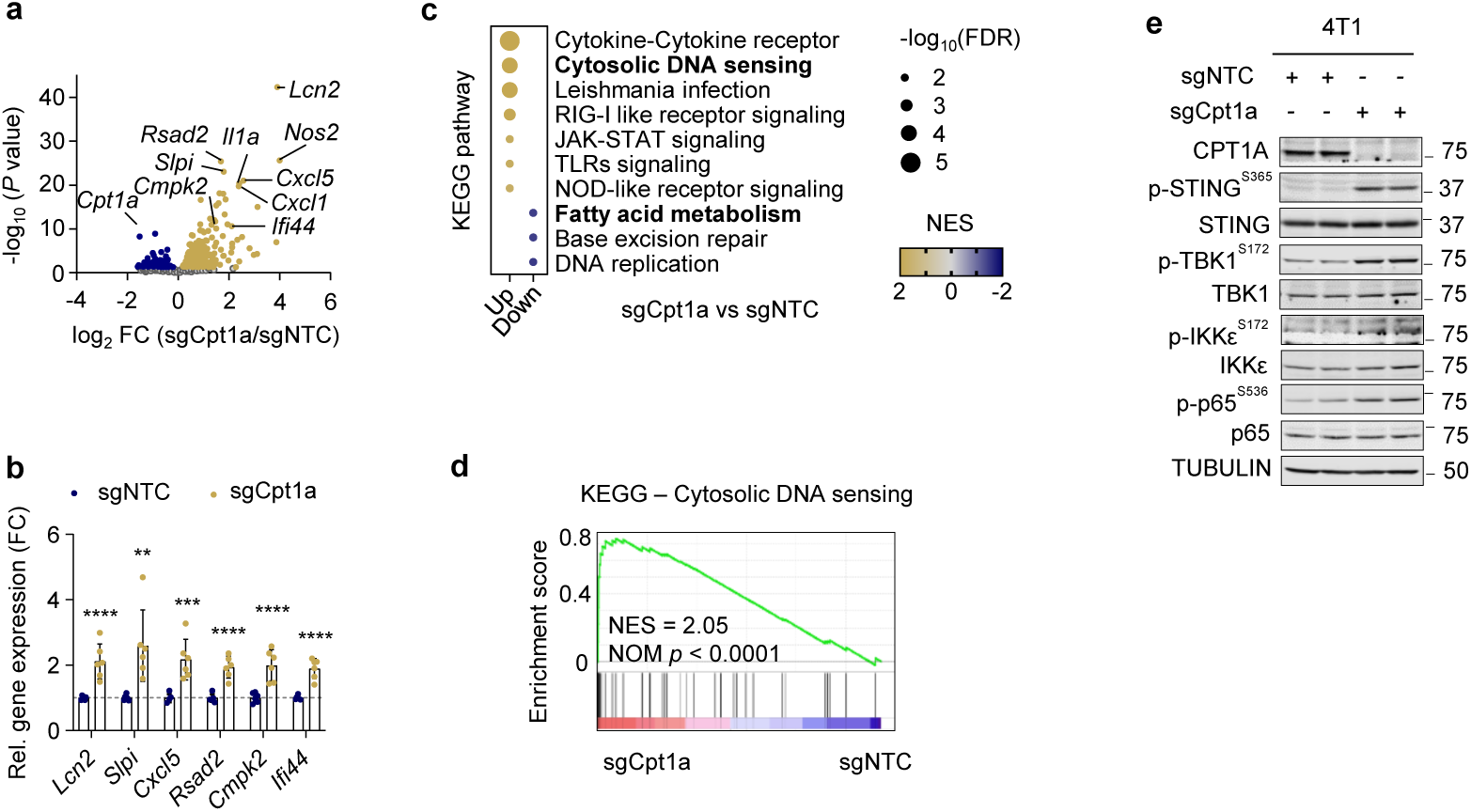
STING chronic activation supports CPT1A-deficient tumor cell metastasis to lung. **a**, Volcano plot displays differential expressed genes in *Cpt1a*-KO (sgCpt1a) 4T1 cells compared to sgNTC controls. **b**, RT-PCR to validate top upregulated genes in (**a**). Unpaired Student’s t-tests were performed. **c**, Bubble plots depicts upregulated (yellow) and downregulated (blue) KEGG pathways in CPT1A-deficient (sgCpt1a) 4T1 cells, relative to controls (sgNTC). FDR, false-discovery rate; NES, normalized enrichment score. **d**, KEGG Cytosolic DNA sensing signatures. **e**, Immunoblot analysis of STING activation in control (sgNTC) and *Cpt1a*-KO (sgCpt1a) 4T1 cells. **p<0.01, ***p<0.005, ****p<0.001. ns, not significant.

Pathway enrichment analysis also revealed enrichment of the cytosolic DNA sensing pathway. The cGAS-Sting-NFκB signaling pathway has emerged as a critical regulator of inflammatory gene expression^28,29^, and indeed, phosphorylation of STING, TBK1, IKK, and p65 subunit of NFkB was increased upon loss of CPT1A, indicating activation of the cytosolic DNA sensing pathway in CPT1A-deficient tumor cells (Fig. 2e).

Consistent with Sting activation, elevated levels of cytosolic DNA were observed in Cpt1a KO cells (Fig. 3a and Extended Data Fig. 3a). While nuclear DNA in the cytoplasm was not affected, cytosolic mitochondrial DNA (mtDNA) was significantly elevated in Cpt1a KO breast cancer cells either by qPCR in cytosol fraction or by immunofluorescence co-staining of antibodies against double strand DNA or Tom20, a mitochondria marker (Fig. 3b-c and Extended Data Fig. 3b-c). Depletion of mtDNA using dideoxycytidine (ddC) reversed the impact of Cpt1a deficiency on cytosolic mtDNA as well as inflammatory gene expression (Fig. 3d-e and Extended Data Fig. 3d-e), indicating that mtDNA release into the cytosol may trigger the pro-inflammatory phenotype in Cpt1a KO cells.

**Fig. 3.**
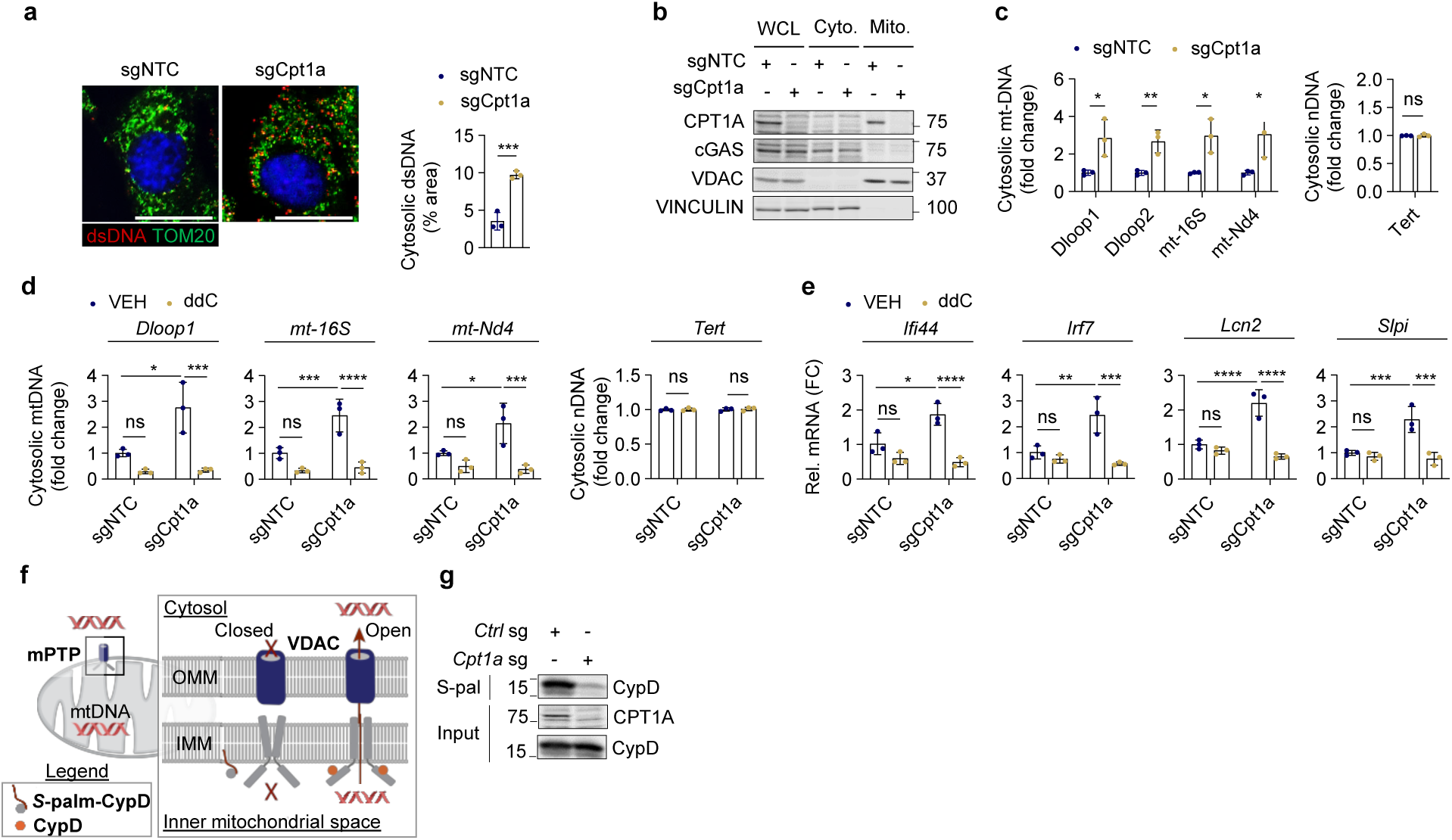
Loss of CPT1A promotes mtDNA release through the mitochondrial permeability transition pore (mPTP). **a**, Immunofluorescence staining of dsDNA (red) and Tom20 (green) in control (sgNTC) and *Cpt1a*-KO (sgCpt1a) 4T1 cells. Scale bar 20µM. Cytosolic dsDNA coverage was determined as a percent (%) of total area. Unpaired Student’s t-test was performed. **b-c**, Fractionation of 4T1 control (Ctrl sg or sgNTC) or *Cpt1a*-KO (Cpt1a sg or sgCpt1a) cells was performed to collect the cytosolic fraction. **b**, Immunoblot to demonstration fractionation. Markers to denote mitochondrial (Mito., CPT1A and VDAC) and cytosolic (Cyto., cGAS and VINCULIN) fractions are shown. Whole cell lysates (WCL) serve as an input control. **c**, Relative levels of mitochondrial DNA (mtDNA) regions (Dloop1, Dloop2, mt-16S, mt-Nd4) (left) or nuclear DNA (nDNA, Tert) (right) in the cytosolic fraction was determined using RT-PCR. Unpaired Student’s t-tests were performed. **d-e**, Control (sgNTC) or *Cpt1a*-KO (sgCpt1a) 4T1 cells were treated with vehicle (VEH) or dideoxycytidine (ddC) to deplete mtDNA. n=3 per group. **d**, Relative levels of mtDNA regions (left) or nDNA (right) in the cytosolic fraction was determined using RT-PCR. Two-way ANOVA with Tukey’s post hoc; p=0.0181 (Dloop1), p=0.0109 (mt-16S), p=0.0312 (mt-Nd4), p=0.886 (Tert). **e**, Relative expression of *Ifi44, Irf7, Lcn2*, and *Slpi* was determined by RT-PCR. Two-way ANOVA with Tukey’s post hoc; p=0.0104 (Ifi44), p=0.00637 (Irf7), p=5.45×10^-4^ (Lcn2), p=0.00357 (Slpi). **f**, Schematic model of mtDNA release from the mitochondrial permeability transition pore (mPTP). CypD acts as regulatory subunit of the mPTP, located within the inner mitochondrial space. VDAC acts as a channel on the outer mitochondrial membrane (OMM). Palmitoylation of CypD at C202 (C203 in humans) prevents CypD from interacting with the mPTP to keep the pore in a closed state. **g**, Immunoblot of S-palmitoylated (S-pal) CypD in control (sgNTC) and *Cpt1a*-KO (sgCpt1a) 4T1 cells. Input shows total CPT1A and CypD levels. *p<0.05, **p<0.01, ***p<0.005, ****p<0.001. ns, not significant.

The mitochondrial permeability transition pore (mPTP) has been shown to release mtNDA independent of apoptotic mechanisms^30–32^ (Fig. 3f). The mPTP is a complex of proteins, including CypD at the inner mitochondrial membrane, with several outer mitochondrial membrane proteins such as VDAC participating in mitochondrial permeabilization^33–35^. Therefore, we assessed the role of the mPTP and VDAC in the pro-inflammatory gene expression in CPT1A KO cells. Knockdown of VDAC/*Vdac1* or CypD/*Ppif* rescues the inflammatory gene expression in Cpt1a KO cancer cells (Extended Data Fig. 3f-g), supporting a role of the mPTP in driving inflammatory response upon Cpt1a loss.

Regulation of the mPTP has been attributed to post-translational modifications of CypD inside the mitochondria, including at the Cys202 site in mice (Cys203 in humans)^35–40^. Palmitoylation of this cysteine residue has been shown to keep the mPTP in a closed state^38–40^. CypD is significantly palmitoylated in 4T1 breast cancer cells which is diminished upon expression of a C202S CypD mutant (Extended Data Fig. 4h). Given CPT1A’s role in transporting long-chain fatty acids, such as palmitate, into the mitochondria, we investigated whether CPT1A could regulate mPTP opening through palmitoylation of CypD (Fig. 3f). Loss of CPT1A dramatically reduces CypD palmitoylation (Fig. 3g), indicating a role of CPT1A in regulating CypD and the mPTP.

### Loss of CPT1A in breast cancer cells creates an immunosuppressive lung metastatic tumor microenvironment and is associated with poor survival in breast cancer patients

Given that the pro-metastatic role of CPT1A loss in breast cancer cells was dependent on lymphocytes (Fig. 1), we evaluated the immune landscape of CPT1A KO lung metastatic tumors. While CD45+ immune cells were unchanged, CPT1A KO lung metastatic tumors shift toward more T cell enriched (Fig. 4a-b). Specifically, CPT1A KO lung metastatic tumors accumulate more immunosuppressive T helper populations, including GATA3+ Th2 and activated FoxP3+ T regulatory T cells (Fig. 4c-e). Fewer multifunctional cytotoxic CD8+ T cells producing IFNγ and TNFα (Fig. 4f) indicate an impaired anti-tumor immune environment in CPT1A KO lung metastatic tumors.

**Fig. 4.**
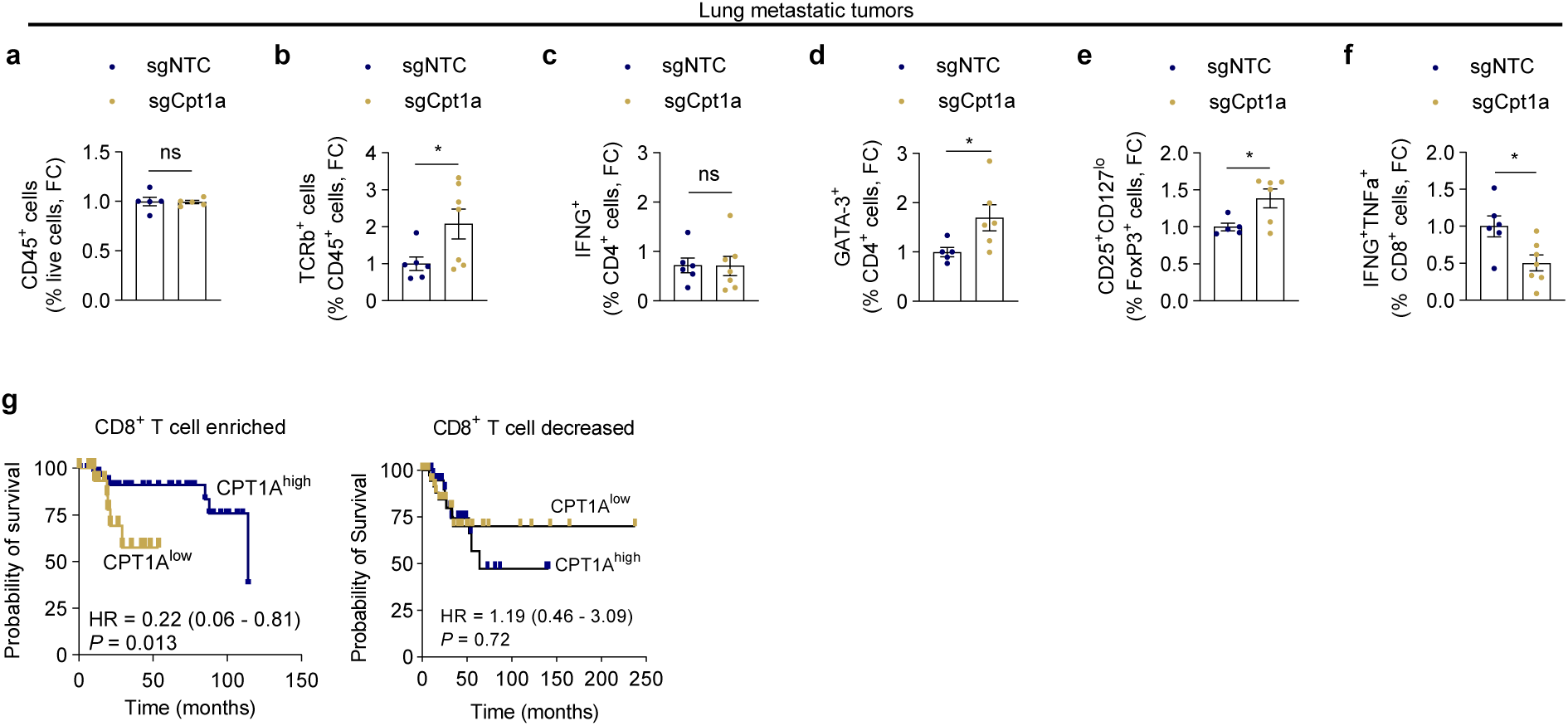
Loss of CPT1A induces immunosuppressive TME in metastatic lesions. **a-f**, The frequencies of the indicated immune cell populations in lung metastatic tumors from female BALB/c WT mice inoculated with control (sgNTC) or *Cpt1a*-KO (sgCpt1a) 4T1 cells was determined by flow cytometry. (n≥5 mice per group). CD45^+^ cells (**a**), TCRβ^+^ T cells (**b**), CD4^+^IFN𝛾^+^ Th1 (**c**), CD4^+^GATA3^+^ Th2 (**d**), CD25+CD127^lo^ activated CD4^+^FoxP3^+^ Treg cells (**e**), and multifunctional cytotoxic IFN𝛾^+^TNFα^+^ CD8^+^ T cells (**f**). Welch’s t-tests were performed. **g**, Overall survival of advanced breast cancer patients stratified by CD8+ T cell tumor enrichment and *CPT1A* expression. Logrank p-values and hazard ratios (HR) are shown with 95% confidence interval range. *p<0.05. ns, not significant.

While low *CPT1A* expression has been previously attributed to improved survival among all breast cancer patients^11,12,17,18,41^, the relationship between tumor cell *CPT1A* expression and immune cells has not been fully explored. Using a gradual model of metastasis that better reflects the patient experience with breast cancer, we show that loss of CPT1A enhances breast cancer metastasis in a T cell dependent manner.

Therefore, we next evaluated the relationship between *CPT1A* and CD8+ T cells on breast cancer patient survival (Fig. 4g). Indeed, we report that low *CPT1A* expression is associated with poorer survival in breast cancer patients with greater enrichment of CD8+ T cells (Fig. 4i). However, the negative impacts of low *CPT1A* expression on survival are lost in patients with poor CD8+ T cell infiltration (Fig. 4i). Overall, these data support an immune-dependent metastatic protective role of CPT1A in breast cancer patients.

## Discussion

The metastatic progression of breast cancer involves complex interactions between tumor cells and immune cells, with T cells playing a key role in anti-tumor immunity^1–3^. While CPT1A is a known intrinsic driver of tumor growth^11–17^, this study utilized an unbiased CRISPR screen to reveal a surprising extrinsic role for CPT1A in immune-mediated suppression of metastasis. Using a gradual metastatic model mimicking patient disease progression, we demonstrate that CPT1A loss enhances lung metastasis in immunocompetent mice through cytosolic mtDNA release via the mitochondrial permeability transition pore (mPTP), leading to a STING-dependent inflammatory response. This inflammatory environment leads to CD8+ T cell dysfunction, impairing anti-tumor immune responses that promotes metastatic outgrowth. Clinically, low CPT1A expression correlates with poorer survival in breast cancer patients with high grade metastatic disease and with high CD8+ T cell infiltration, underscoring the immune-dependent metastatic protective function of CPT1A. These findings advance our mechanistic understanding of how metabolism influences immune cell responses during metastatic progression in breast cancer.

It is becoming increasingly clear that metastatic progression is driven by interactions between tumor and immune cells^1–5^. While CPT1A has been well studied as an intrinsic driver of metastasis^11–17^, we show that CPT1A also has a dramatic extrinsic influence to suppress metastasis. The extensive use of immunodeficient mouse metastasis models has limited our understanding of how CPT1A impacts tumor-immune interactions^11–16^.

Similar tumor cell intrinsic anti-metastatic impacts of CPT1A deficiency have been described in an immunocompetent host^17^. However, immunosuppression has been shown to build gradually during metastatic progression^23,24^, indicating the importance of using extended metastasis models to better capture the full interaction between metastatic cancer cells and immune cells. Using a metastasis model that better recapitulates breast cancer progression in patients, our data reveal that CPT1A loss drives metastasis in a lymphocyte-dependent manner. A pro-metastatic role of chromosomal instability-derived inflammation has been described in similar gradual progressive metastasis models^42,43^. These results emphasize the importance of studying metastasis in models that account for the full reciprocity of immune-tumor dynamics, as metabolic enzymes such as CPT1A may influence immune function in a gradual, progressive manner.

Beyond its role in fatty acid oxidation, our data indicates that CPT1A’s catalytic activity protects against inflammation within breast cancer cells. CPT1A loss drives inflammatory gene expression in breast cancer cells via mtDNA release through the mitochondrial permeability transition pore (mPTP). Regulation of mPTP opening occurs inside the mitochondrial matrix through post-translational modifications^35–40^.

Palmitoylation of CypD, the inner mitochondrial regulatory mPTP component, at Cys202 (Cys203 in humans) has been shown to keep the mPTP closed^38–40^. Indeed, we show that CPT1A loss decreases CypD palmitoylation. In the absence of CPT1A, less palmitate is transported into the mitochondria to reduce fatty acid oxidation, thus limiting intramitochondrial palmitate as a modification substrate. Strikingly, a catalytically-dead CPT1A mutant maintains the metastastic phenotype associated with CPT1A loss, demonstrating the critical protective nature of CPT1A’s enzymatic function.

STING activation has been explored as a strategy to enhance anti-tumor responses^44–47^, but STING agonists have shown mixed results in clinical trials^48,49^. Our study reveals that mtDNA release upon CPT1A loss supports STING activation, leading to enhanced inflammation and metastatic progression, a mechanism that aligns with prior findings indicating pro-metastatic inflammation associated with nuclear DNA release^42,43^. These insights suggest that inhibition of STING signaling may be an effective therapeutic strategy for some patient subsets with metastatic breast cancer. In particular, our study reveals that those with low *CPT1A* expression and high CD8+ T cell infiltration may benefit from STING inhibition by mitigating pro-metastatic inflammation, offering new therapeutic options to metastatic breast cancer.

## Methods

### Cell lines

4T1 murine breast cancer cells and human epithelial-like HEK293T cells were purchased from ATCC. E0771 cells were provided by Barbara Fingleton (Vanderbilt University), and MMTV-PyMT were isolated from primary murine C57BL/6 mammary tumors^50^ and provided by Rebecca Cook (Vanderbilt University). All cell lines were cultured in DMEM supplemented with 10% FBS and 100U/mL penicillin-streptomycin, as previously described^51–55^. Routine mycoplasma testing was performed (Invivogen #rep-mys-10).

To generate knockout cell lines, independent sgRNA sequences targeting Cpt1a (sgCpt1a) were cloned into lentiCRISPRv2 puro, a gift from Brett Stringer (Addgene #98290). For CypD (sgPpif) knockout, CRISPR ribonucleoprotein complexes consisting of recombinant Cas9 protein and synthetic sgRNAs were delivered into cells using Lipofectamine CRISPRMAX Cas9 Transfection Reagent (Thermo Fisher Scientific, Cat# CMAX00008) according to the manufacturer’s instructions. A non-targeting sgRNA (sgNTC) was used as a control. Sequences are shown in Supplementary Table 1. To minimize potential off-target effects, at least two independent sgRNAs were evaluated for each gene, and consistent phenotypes were confirmed across sgRNAs. For sgPpif, knockout clones were validated and combined to generate a pooled population. All functional assays were performed using cell populations generated with Cpt1a_sg1 or Ppif_sg_2, as indicated.

For rescue experiments, murine wild-type Cpt1a (WT) or the catalytic mutant CPT1A-D454G (GAC to GGC) open-reading frame sequences flanked by XbaI and Sal1 sites were cloned into pLentiCMV-Blast (Addgene #125133). Mutant constructs were generated by site-directed mutagenesis using the Q5 Site-Directed Mutagenesis Kit (New England Biolabs #E0554) according to the manufacturer’s instructions. CRISPR-resistant constructs were generated by introducing silent mutations within the PAM motif^56^. Re-expression and knockout efficiencies were validated by immunoblotting and genomic DNA sequencing. Re-expression levels were comparable to endogenous protein levels in parental cells.

### Mouse strains

All mouse strains were purchased from The Jackson Laboratory. To generate Cas9-tolerant mice for 4T1 spontaneous metastasis models, Cas9-C57BL/6 (#000664) mice were crossed with BALB/c wild-type (WT) mice (#000651), and the F1 progeny (Cas9^F^^1^) were used for experiments. *In vivo* CRISPR screens and 4T1 spontaneous breast cancer metastasis assays were performed using 6-8-week-old female BALB/c-Rag1^-/-^(#033531), BALB/c WT, or Cas9^F^^1^ mice, as indicated. Syngeneic C57BL/6J WT mice (#000664), Cas9-C57BL/6, or C57BL/6-Rag1^-/-^ (#034159) mice were used for the E0771 metastasis models, as indicated.

### *In vivo* CRISPR screening

A metabolism-focused loss-of-function sgRNA library was utilized, targeting 101 metabolic genes with four sgRNAs per gene as well as ten non-targeting control sgRNAs (Supplemental Table 2). sgRNA oligonucleotides with flanking adaptor sequences (5′-TATCTTGTGGAAAGGACGAAACACCG-[20 bp sgRNA sequence]-GTTTTAGAGCTAGAAATAGCAAGTTAAAAT-3′) were synthesized as a pooled library (Twist Bioscience) and cloned into lenti-gRNA-puro (Addgene #84752) following published protocols^57^ with minor modifications. Briefly, adaptor sequences were appended by PCR amplification using Herculase II Fusion DNA Polymerase (Agilent #600675), followed by gel purification (QIAquick Gel Extraction Kit, Qiagen #28704). Amplified products were assembled into linearized lenti-gRNA-puro using Gibson Assembly Master Mix (New England Biolabs #E2611L). The plasmid library was electroporated into ElectroMAX DH10B cells (Thermo Fisher #18290015) and plated on ampicillin agar plates to achieve >50-fold representation per sgRNA. Colonies were pooled, and plasmid DNA was isolated using an EndoFree Plasmid Maxiprep Kit (Qiagen, Cat# 12362).

To establish Cas9-expressing cells, breast cancer cells were transduced with lenti-Cas9-blast (a gift from Feng Zhang, Addgene #52962) lentivirus. Cas9 expression was validated prior to downstream genome editing experiments. Cas9-expressing 4T1 cells were transduced with serial dilutions of viral supernatant. MOI was maintained <0.3 to favor single sgRNA integration per cell. Transduction efficiency was calculated based on puromycin survival, and only populations meeting MOI criteria were used for downstream screening.

For in vivo screening, 6–8-week-old female BALB/c WT (strain# 000651) or BALB/c-Rag1^-/-^ mice were used. Library-transduced 4T1-Cas9 cells were injected bilaterally into the fourth mammary fat pads. For each injection site, 2.5 × 10^5^ cells representing ∼1,000× sgRNA coverage were suspended in PBS and mixed 1:1 with Matrigel. An aliquot of the transduced cell pool was cryopreserved at implantation and served as the baseline reference (input) for sgRNA distribution analysis. Tumor growth was monitored and resected at day 21 or when average tumor volume reached ∼500 mm^3^. Resected tumors were snap-frozen in liquid nitrogen for analysis. Mice were euthanized upon reaching humane endpoints, and whole lungs were harvested and processed for sgRNA abundance profiling.

### Genomic DNA extraction from cells and mouse tissues

Genomic DNA from cultured cells was isolated using the PureLink Genomic DNA Mini Kit (Thermo Fisher Scientific #K182001) according to the manufacturer’s instructions. For mouse tissues, genomic DNA was extracted using a modified high-yield salting-out protocol adapted from prior studies^58^. Briefly, 100–200 mg fresh or frozen tissue was minced and incubated in 6 mL NK lysis buffer (50 mM Tris-HCl, 50 mM EDTA, 1% SDS, pH 8.0) supplemented with 30 μL Proteinase K (20 mg/mL; Thermo Fisher Scientific #25530049) overnight at 55°C with gentle agitation. RNase A (30 μL, 10 mg/mL; Invitrogen #EN0531) was added the next day and incubated at 37°C for 30 min.

Proteins were precipitated by adding 2 mL pre-chilled 7.5 M ammonium acetate (Millipore Sigma #A2706), followed by vortexing and centrifugation at 4,000 × g for 10 min. The clarified supernatant was transferred to a new tube and DNA was precipitated with an equal volume of 100% isopropanol (Millipore Sigma #I9030). DNA pellets were washed with 70% ethanol, air-dried, and resuspended in 500 μL TE buffer (Thermo Fisher #J75893). Samples were incubated at 65°C for 1 h and further dissolved overnight at room temperature. DNA concentration and purity were determined by spectrophotometry.

### Amplification of sgRNA cassettes for deep sequencing

#### PCR#1: sgRNA cassette amplification

sgRNA representation in plasmid libraries and genomic DNA samples was determined using a two-step PCR strategy optimized to preserve library complexity. The sgRNA cassette was amplified using vector-specific primers (LenIn-Fwd: 5’-AATGGACTATCATATGCTTACCGTAACTTGAAAGTATTTCG-3’; LenIn-Rev: 5’-TCTACTATTCTTTCCCCTGCACTGT-3’). PCR reactions were performed using KAPA HiFi HotStart ReadyMix (Roche #501965217) using the following cycling conditions: 94°C for 5 min; 15–30 cycles of 94°C for 15 s, 65°C gradient annealing (−1°C per cycle) for 30 s, and 72°C for 40 s; followed by 72°C for 5 min. Template inputs were scaled by sample type, including plasmid DNA (100ng), genomic DNA from input cells (1 μg), and genomic DNA from tumor samples (5 μg). Multiple parallel PCR#1 reactions were performed per biological sample to preserve library complexity, and products from the same sample were pooled prior to indexing.

#### PCR#2: indexing and library construction

Pooled PCR#1 products were subjected to a second PCR using custom indexing primers containing Illumina P5 and P7 adapter sequences and sample-specific index barcodes (Supplementary Table S3). For each biological sample, a minimum of two 100 μL PCR#2 reactions were performed, using 10 μL pooled PCR#1 product per reaction (∼1 reaction per 10^4^ constructs). PCR#2 products were pooled and normalized prior to combining uniquely indexed samples. Libraries were size-selected on 2–3% agarose gels and purified using the QIAquick Gel Extraction Kit (Qiagen #28704). Final library concentrations were determined using the Qubit dsDNA High-Sensitivity assay (Invitrogen #Q32851) and gel-based quantification. Libraries were sequenced in paired-end mode (150 bp) on the Illumina MiSeq platform at the Vanderbilt Technologies for Advanced Genomics core, with 5–20% PhiX spike-in to improve sequence diversity.

Throughout library preparation and sequencing, ≥1,000-fold sgRNA representation was maintained.

### CRISPR screen analysis

Raw FASTQ files were processed using MAGeCK (v0.5.0.3). Reads were trimmed and aligned to the reference sgRNA library to generate count matrices. Read counts were median-normalized across samples, and differential sgRNA abundance was calculated using a negative binomial model. sgRNA enrichment/depletion was evaluated relative to non-targeting controls, and gene-level scores were derived from aggregated sgRNA statistics, using default MAGeCK parameters.

### Gene silencing

For transient gene silencing experiments, murine breast cancer cells were transfected with ON-TARGETplus SMARTpool siRNAs (25 nM) obtained from Dharmacon (Horizon Discovery) targeting mouse *Cpt1a* (Cat# L-042456-01-0010), *Tmem173*/STING (Cat# L-055528-00-0010), *Ppif*/CypD (Cat# L-062722-01-0010), *Vdac1* (Cat# L-047345-00-0010), and a scrambled non-targeting control (Cat# D-001810-10-20) using Lipofectamine RNAiMAX (Invitrogen, Cat# 13778150) according to the manufacturer’s instructions. Knockdown efficiency was verified at the mRNA and/or protein level by qRT-PCR or immunoblotting prior to functional assays.

### Gradual spontaneous syngeneic metastasis models

In a gradual spontaneous orthotopic model of metastasis, 2.5×10^5^ 4T1 or E0771 cells were implanted into the inguinal mammary fat pad of 6-to-8-week old female syngeneic mice. To enable spontaneous metastatic dissemination, primary tumors were surgically resected at 14-21 days and snap frozen for further analysis. Tumor dimensions were measured using digital calipers, and tumor volume was calculated as: Volume = L × (W/2)^2^, where L represents tumor length and W represents tumor width. Lungs were harvested 3-8 weeks after resection. Lung metastases were quantified by one or more of the following methods: (1) enumeration of macroscopic metastatic nodules, (2) qPCR targeting the Cas9 vector cassette, and/or (3) deep sequencing–based quantification of sgRNA.

For competition metastasis assays, Cpt1a KO and non-targeting control cells derived from 4T1 or E0771 lines were mixed at a 1:1 ratio prior to implantation. 2.5×10^5^ mixed cells were injected into the inguinal mammary fat pad of 6-to-8-week-old female syngeneic mice. An aliquot of the injected cell mixture was cryopreserved as the baseline input reference for downstream sgRNA quantification. Tumors were resected at day 14-18, and primary tumors were snap-frozen in liquid nitrogen for genomic DNA extraction. To assess spontaneous metastatic dissemination, lungs were harvested 2-4 weeks post-resection. Tissues were mechanically dissociated on ice and genomic DNA was extracted as described above. Genomic DNA from input mixtures, primary tumors, and lungs was subjected to NGS-PCR and deep sequencing. Relative sgRNA abundance was quantified using MAGeCK. sgRNA counts in tumors and metastatic lesions were normalized to the corresponding sgNTC abundance and referenced to the input population to calculate relative depletion or enrichment.

### Metabolite profiling

Cell pellets of control (sgNTC) or Cpt1a-KO (sgCpt1a) 4T1 cells (n=6 per sample) were analyzed by Metabolon using ultra-high performance liquid chromatography-tandem mass spectrometry (UPLC-MS/MS), as previously described^54^. Compounds were identified based on retention index, library mass matching, and score matching with authentic standards. 837 metabolites were identified, and abundance was normalized to sample protein levels. Scaled results were grouped based on major classes and subclasses.

### Long-chain fatty acid beta-oxidation stress assays

The Seahorse XF Palmitate Oxidation Stress Test Kit (Agilent #103693-100) was performed according to manufacturer’s instructions to assess long-chain fatty acid beta-oxidation in Cpt1a-deficient cells. 5×10^4^ cells control (sgNTC) or Cpt1a-KO (sgCpt1a) 4T1, E0771, or PyMT cells were seeded in Seahorse XFe24 microplates (Agilent, Cat# 101122-100) in complete growth medium. After 24 h, cells were switched to palmitate-limited medium consisting of XF DMEM supplemented with 0.5 mM glucose, 1 mM glutamine, and 150 μM L-carnitine and incubated overnight at 37°C. Cells were then washed and provided with palmitate oxidation stress test assay medium containing 1 mM glucose, 0.5 mM L-carnitine, 170 μM palmitate, and 0.7 mM glutamine. Oxygen consumption rates (OCR) were measured using a Seahorse XFe24 Flux Analyzer, with rates normalized to total protein, as determined by the Bio-Rad DC Protein Assay Kit (Bio-Rad, Cat# 5000112).

### Cell viability assay

Cell viability was assessed using an MTT-based assay. Briefly, 1×10^4^ control (sgNTC) or Cpt1a-KO (sgCpt1a) 4T1, E0771, or PyMT cells were seeded in 96-well plates. Viability was determined using the CyQUANT MTT Cell Viability Assay Kit (Thermo Fisher Scientific #V13154) at the indicated time points according to the manufacturer’s instructions. Absorbance (570 nm) was determined using a BioTek Synergy H1 microplate reader. Background-subtracted absorbance values were used as a surrogate for viable cell number.

### Lung histology

Right lung lobes were harvested and fixed in 4% paraformaldehyde overnight at 4°C, then transferred to 70% ethanol. Paraffin embedding, sectioning, and hematoxylin and eosin (H&E) staining were performed by the Vanderbilt University Medical Center Translational Pathology Shared Resource.

### RNA sequencing

For RNA sequencing of cultured cells, RNA was extracted from control (sgNTC) or Cpt1a-KO (sgCpt1a) 4T1 cells 4T1-sgCpt1a and 4T1-sgNTC cells using the RNeasy Plus Mini Kit (Qiagen #74134) according to the manufacturer’s protocol. RNA quantity and quality were assessed using the Qubit High-Sensitivity RNA assay (Invitrogen #Q32852). RNA sequencing was performed by Azenta using the Illumina Novaseq platform. Sequencing reads were trimmed and mapped to the mouse reference genome GRCm38 using STAR aligner (v2.7.9a). Gene-level counts were generated using the SummarizeOverlaps function from the GenomicAlignments R package. Differential gene expression analysis between groups was performed using DESeq2 (v1.24.0). Adjusted p values were calculated using the Benjamini–Hochberg method to control the false discovery rate (FDR). Log2 fold change (FC) values were computed relative to controls.

Pathway enrichment analyses were performed using Gene Set Enrichment Analysis (GSEA) software (Broad Institute, v4.4.0). The KEGG_LEGACY pathway database was queried after mapping mouse genes to human orthologs using the biomaRt (v2.52.0) in R (v4.2.1). Enrichment results were evaluated using normalized enrichment scores (NES) and false discovery rate (FDR) q-values calculated by permutation testing within GSEA.

### Quantitative real-time PCR

To evaluate mRNA expression, total RNA was isolated from cells using the RNeasy Plus Mini Kit (Qiagen #74134). 1μg of total RNA was reverse-transcribed using the iScript cDNA Synthesis Kit (Bio-Rad #1708891) according to manufacturer’s instructions. Using the sequences defined in Extended Data Table 4, quantitative real-time PCR (RT-PCR) was performed at least in duplicate using iTaq Universal SYBR Green Supermix (Bio-Rad #1725121) on the Quant Studio 3 (Applied Biosystems). Relative gene expression was calculated using the ΔΔCt method.

### Subcellular fractionation for mtDNA release assays

To assess mitochondrial DNA (mtDNA) release into the cytosol, subcellular fractionation was performed using a modified digitonin-based permeabilization protocol adapted from published methods^59^. Briefly, cytosolic and mitochondrial-enriched fractions were obtained by incubating the cell pellet in digitonin lysis buffer (50 mM HEPES, pH 7.4; 150 mM NaCl; 200-400 μg/mL digitonin; supplemented with protease inhibitors) to selectively permeabilize the plasma membrane while preserving mitochondrial and nuclear integrity. The cytosolic fraction was obtained from the clarified supernatant. The remaining pellet was washed and resuspended in NP-40 lysis buffer (Research Products International #50-197-8071) supplemented with protease inhibitors, incubated on ice for 10 min, and centrifuged at ∼21,000 × g for 10 min at 4°C to generate the mitochondria-enriched fraction. Fraction purity was validated by immunoblotting using cytosolic markers (cGAS, Vinculin) and mitochondrial markers (CPT1A, VDAC). Whole-cell extracts (WCE) were obtained as input controls.

For mitochondrial DNA (mtDNA) quantification, DNA was isolated from whole-cell lysates, cytosolic fractions, or mitochondria-enriched fractions using the QIAamp DNA Mini Kit (Qiagen #51304). Multiple regions (*Dloop1, Dloop2, mt-16S, mt-Nd4*) of the murine mitochondrial genome (NC_005089.1) were amplified in the cytosolic fraction (Supplementary Table 4). The nuclear gene *Tert* was used as a nuclear DNA (nDNA) control. mtDNA abundance was normalized to nuclear Tert levels using the ΔΔCt method.

### Immunoblotting

Cells were lysed in RIPA buffer (Sigma-Aldrich #R0278) supplemented with protease and phosphatase inhibitors for 30 min on ice, followed by centrifugation to remove debris. Cytosolic and mitochondria-enriched fractions were prepared as described above. Protein concentrations were determined using the DC Protein Assay Kit II (Bio-Rad #5000112). Equal amounts of protein were resolved by SDS-PAGE and transferred to 0.2 μm nitrocellulose membranes. Membranes were blocked in INTERCEPT TBS Blocking Buffer (LI-COR Biosciences #927-60010) and incubated with the following primary antibodies at 1:1000, unless indicated: Cas9 (Cell Signaling Technology (CST) #14697), CPT1A (Abcam #ab128568 or Abcam #ab234111 or Proteintech #15184-1-AP), phospho-STING (Ser365, CST #72971), STING (CST #50494), phospho-TBK1 (Ser172, CST #5483), TBK1 (CST #3504), phoshpo-IKKε (Ser172, Invitrogen #PA5-105888), IKKε (CST #3416), phospho-p65 (Ser536, CST #3033), p65 (CST #8242), CypD (Thermo Fisher #455900), cGAS (CST #31659), VDAC (XXX), Vinculin (CST #13901), or β-Tubulin (Sigma #T4026). Subsequently, membranes were incubated with IRDye 680LT goat anti-mouse (1:20,000; LI-COR #926-68020), IRDye 680LT goat anti-rabbit (1:20,000; LI-COR #926-68021), IRDye 800CW goat anti-mouse (1:10,000; LI-COR #926-32210), or IRDye 800CW goat anti-rabbit (1:10,000; LI-COR #926-32211).

Proteins were detected using a LI-COR Odyssey imaging system.

### Immunofluorescence

Cells were seeded on #1.5 glass coverslips (VWR, Cat# 48393-241) and fixed in 4% paraformaldehyde for 30 min, permeabilized with 0.1% Triton X-100 for 5 min, and blocked in 5% BSA in PBST. Cells were incubated overnight at 4°C with mouse anti-dsDNA (1:500; Millipore #CBL186) and rabbit anti-TOM20 (1:400; CST #42406). After washing, cells were incubated with Alexa Fluor 488 goat anti-mouse IgG (1:1000; Thermo Fisher Scientific #A-11001) and Alexa Fluor 594 goat anti-rabbit IgG (1:1000; Thermo Fisher Scientific #A-11012) for 1 h at room temperature. Nuclei were counterstained with DAPI (Thermo Fisher Scientific #D1306) and coverslips were mounted using ProLong Gold Antifade Mountant (Molecular Probes #P36971). Images were acquired using an Olympus BX60 compound microscope equipped with a 60× oil immersion objective. Image analysis was performed using FIJI/ImageJ (v1.48).

### Mitochondrial DNA (mtDNA) depletion

Cells were cultured in complete growth medium supplemented with 2′,3′-dideoxycytidine (ddC; 100 nM) for 21 days to deplete mtDNA. Fresh ddC-containing medium was replaced every 48 h. Vehicle-treated cells cultured in parallel served as controls. mtDNA depletion was validated by efficient loss of cytosolic mtDNA depletion.

### CypD S-palmitoylation

S-palmitoylation of CypD was assessed using the CAPTUREome S-Palmitoylated Protein Kit (Badrilla #K010-311) according to the manufacturer’s protocol. Briefly, cells were conditioned in palmitate-free medium overnight prior to addition of 50 μM palmitate-BSA. After 24 hours, mitochondria-enriched fractions were prepared from harvested cells by differential centrifugation, as described above, and equal protein was subjected to thiol blocking, hydroxylamine-dependent thioester cleavage (or cleavage control), and resin capture. Bound proteins were eluted and analyzed by immunoblotting, as described above.

### Flow cytometry

Resected primary tumors and tumor-enriched lung metastatic tissues were dissociated in RPMI-1640 containing 5% FBS, collagenase IA (1 mg/mL; Sigma-Aldrich #C9891), and DNase I (0.25 mg/mL; Sigma-Aldrich #DN25), as previously described^51,54,55^. Cells were filtered through a 70 μm cell strainer and subjected to red blood cell lysis. Cells were stained with the Ghost Violet 510 viability dye (Biotek Bioscience #13-0870) and blocked using mouse anti-CD16/32 (Cytek Bioscience #70-0161). Extracellular staining was performed, using antibodies against CD45, TCRβ, CD4, CD8, CD11b, CD25, and/or CD127 (see Supplementary Table 5). To detect TNFα and IFNγ, the Cytofix/Cytoperm kit (BD Biosciences #554714) was used according to manufacturer’s instructions. For detection of the GATA3 and FoxP3 transcription factors, the FoxP3/Transcription Factors Staining Kit (Cytek Biosciences #TNB-0607) was used as directed. Appropriate compensation controls were used, along with fluorescence minus one (FMO) and isotype controls, where appropriate. Flow cytometry was performed on a BD Fortessa using BD FACS Diva software. Analysis was completed using FlowJo (v10). Data were normalized to controls and shown as fold change (FC).

### Survival analysis

Overall survival data in breast cancer were downloaded from KM plot (www.kmplot.com)^60^. Analyses were restricted to primary tumors from patients with a with available overall survival data. Patients with advanced breast cancer, defined as Grade 3 with lymph node involvement. Patients were stratified based on low (quartile 1, Q1) or high (Q2-Q4) *CPT1A* expression from available RNA-seq data and restricted to either enriched or decreased CD8+ T cell content, as indicated.

### Statistics

All statistical analyses were performed using GraphPad Prism (v9.0 or later) and R (v4.0 or later) unless otherwise indicated. Data are presented as mean ± SEM. Comparisons between two groups were performed using unpaired two-tailed Student’s t-tests or Welch’s t-test, as indicated. For multiple group comparisons, one-way or two-way analysis of variance (ANOVA) with appropriate post hoc correction was applied.

Outliers were excluded based on the ROUT method (5%). Statistical differences with a p-value less than 0.05 were considered to be statistically significant. Sample sizes (n) represent independent biological replicates as defined in the figure legends. Investigators were blinded to group allocation during histological quantification where feasible.

For survival studies using the 4T1 spontaneous metastasis model, group sizes of 15 mice were determined based on prior power calculations, providing 80% power at 95% confidence to detect hazard ratios below 0.25 or above 4.0. Survival curves were analyzed using the Kaplan–Meier method and compared using the Mantel–Cox log-rank or Gehan-Breslow-Wilcoxon test, as indicated. Hazard ratios (HR) and 95% confidence intervals were determined using the log-rank method. Bonferroni’s correction was applied for multiple comparisons.

### Study approval

All experiments involving animals were approved by the Institutional Animal Care and Use Committee (IACUC) at Vanderbilt University Medical Center (Nashville, TN).

## Acknowledgements

The authors would like to thank Jeff Rathmell (University of Chicago) for assistance with performing the CRISPR screen. This work was supported by a Career Enhancement Award to DNE, funded through the Vanderbilt-Ingram Cancer Center SPORE in Breast Cancer (P50CA098131). DNE is also supported by a Department of Defense CDMRP grant W81XWH2210109. JC is supported by NIH grants CA250506 and CA271176, a VA Career Scientist Award (5IK6BX005391), and a VA Merit Award (5101BX000134).

Flow cytometry experiments were performed in the Vanderbilt University Medical Center (VUMC) Flow Cytometry Shared Resource, supported by the Vanderbilt-Ingram Cancer Center (P30 CA68485) and the Vanderbilt Digestive Disease Research Center (DK058404). Tissue processing was performed by the VUMC Translational Pathology Shared Resource, which is supported by a NCI/NIH Cancer Center Support Grant (P30CA068485). Next Generation Sequencing and digital spatial profiling was performed by the Vanderbilt Technologies for Advanced Genomics (VANTAGE) Core.

