## Supplementary Materials for "CPT1A loss promotes lung metastasis in immune-competent mice via a mechanism of mtDNA release and chronic activation of STING pathway"

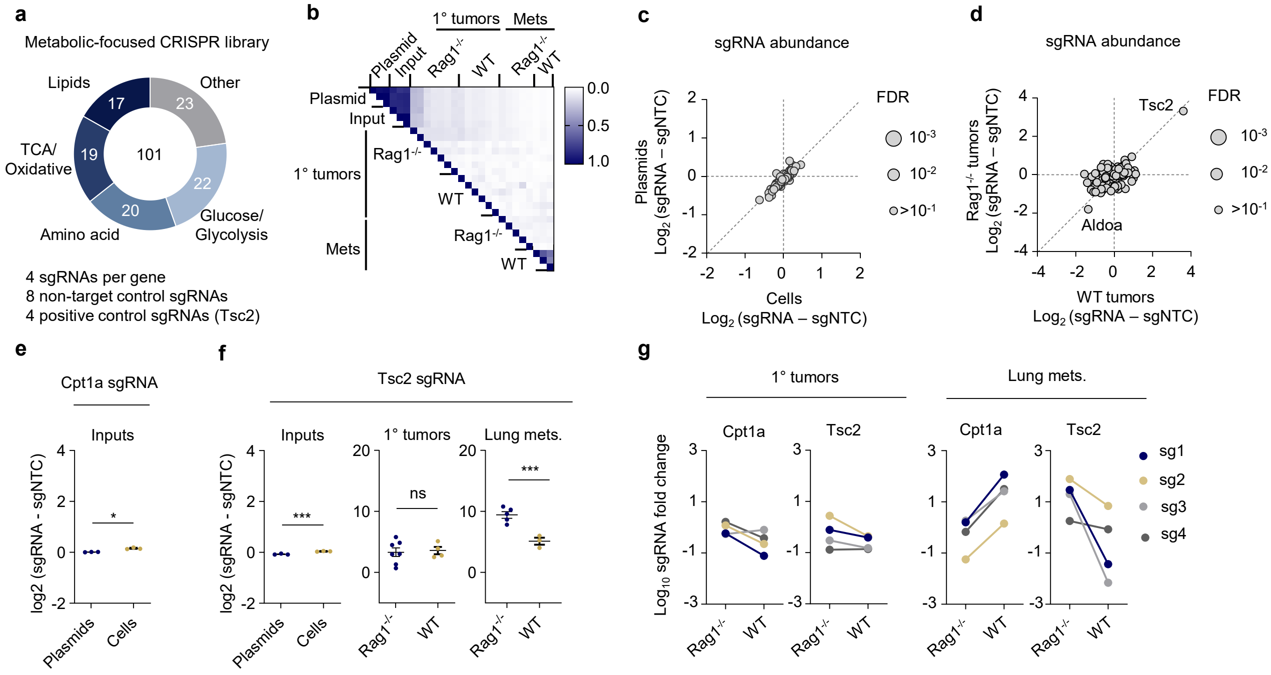

**Extended Data.1 | Metabolic-focused *in vivo* CRISPR KO screen identifies CPT1A as an immune-dependent suppressor of metastasis.** **a**, Distribution of genes in the metabolic-focused CRISPR KO library. **b**, Pearson correlation coefficient of the normalized sgRNA read counts across plasmid, input cells when performed orthotopic transplantation (day3 after antibiotic selection), primary tumors (1° tumors, 3 weeks after transplantation) and whole lung with outgrowth metastatic tumors (Lung mets., 4-9 weeks after tumor resection) from both Rag1$^{-/-}$ and BALB/c WT mice. $n \geq 3$ mice per group. **c-d**, The differential abundance of sgRNA corresponding genes between plasmid library and input cells (**c**) or in primary tumors between Rag1$^{-/-}$ and BALB/c WT mice (**d**). Target gene sgRNAs were normalized to non-targeted control sgRNA (sgNTC). **e-f**, Normalized log2-transformed read counts of *Cpt1a* in input cells (**e**) or the tumor growth suppressor *Tsc2* in input cells, primary tumors, or lung metastases (**f**). $n \geq 3$ mice per group. Welch's t-tests were performed. **g**, $\text{Log}_{10}$ transformed read counts of individual Cpt1a or Tsc2 sgRNAs in primary tumors and lung metastases from Rag1$^{-/-}$ (n=7) or BALB/c (n=4) mice. *p<0.05, ***p<0.005. ns, not significant.

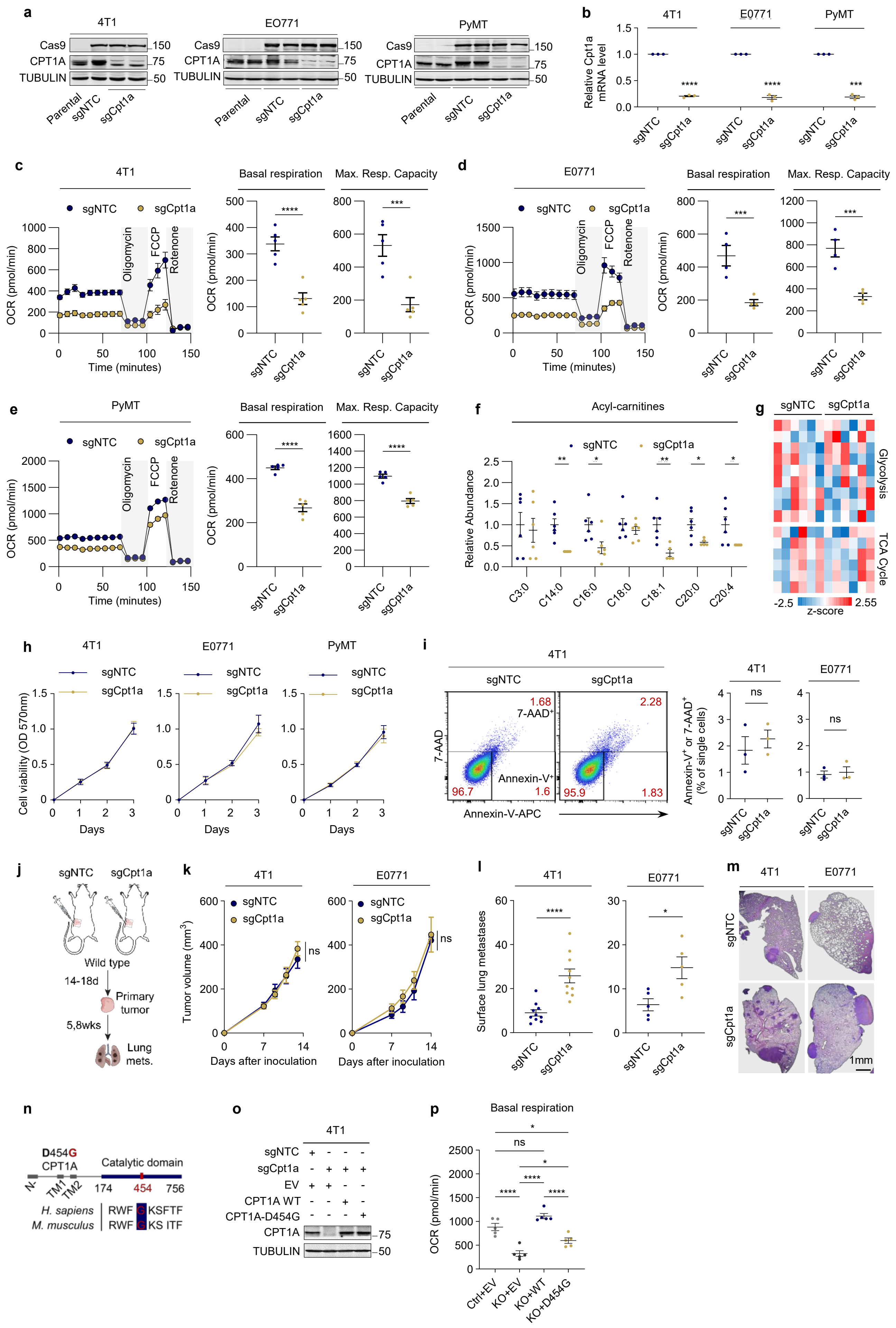

a
4T1
E0771
PyMT
Cas9
CPT1A
TUBULIN
Parental
sgNTC
sgCpt1a
b
Relative Cpt1a mRNA level
c
4T1
sgNTC
sgCpt1a
OCR (pmol/min)
Oligomycin
FCCP
Rotenone
Time (minutes)
Basal respiration
Max. Resp. Capacity
d
E0771
e
PyMT
f
Acyl-carnitines
Relative Abundance
C3:0
C14:0
C16:0
C18:0
C18:1
C20:0
C20:4
g
Glycolysis
TCA Cycle
z-score
h
Cell viability (OD 570nm)
Days
i
7-AAD
Annexin-V-APC
7-AAD+
Annexin-V+
96.7
1.68
1.6
95.9
2.28
1.83
Annexin-V+ or 7-AAD+ (% of single cells)
ns
j
Wild type
14-18d
Primary tumor
5,8wks
Lung mets.
k
Tumor volume (mm3)
Days after inoculation
l
Surface lung metastases
m
1mm
n
D454G
CPT1A
Catalytic domain
N-
TM1
TM2
174
454
756
H. sapiens
M. musculus
RWF G KSFTF
RWF G KS ITF
o
EV
CPT1A WT
CPT1A-D454G
p
Basal respiration
Ctrl+EV
KO+EV
KO+WT
KO+D454G

**Extended Data.2 | CPT1A deficiency impairs cellular respiration and increases metastatic tumor volume in the lung. a**, Immunoblots of CPT1A and Cas9 in parental, sgNTC control and *Cpt1a*-KO (sgCpt1a) in 4T1, EO771 and PyMT (BL6). **b**, Relative *Cpt1a* mRNA expression (fold change) determined by qPCR in control (sgNTC) and CPT1A-deficient (sgCpt1a) 4T1, EO771 and PyMT cells. Unpaired Student's t-tests were performed. **c-e**, Oxygen consumption rate (pmol/min) of control (sgNTC) or CPT1A-deficient (sgCpt1a) 4T1 (**c**), EO771 (**d**), or PyMT (**e**) cells was determined (per $5 \times 10^4$ cells) (n=4-5) using the Seahorse long-chain fatty acid oxidative stress assay kit. Basal respiration and maximum respiration capacity OCR data are shown. Unpaired Student's t-test was performed. **f**, Relative abundance of acyl-carnitines in control (sgNTC) or *Cpt1a*-KO (sgCpt1a) 4T1 cells. Welch's t-tests were performed. **g**,Heat map showing relative abundance of metabolites associated with glycolysis or TCA cycle in control (sgNTC) or *Cpt1a*-KO (sgCpt1a) 4T1 cells. Welch's t-tests indicated no differences in metabolite abundance between groups. **h**, Control (sgNTC) or Cpt1a-deficient (sgCpt1a) 4T1, EO771 and PyMT tumor cell viability at indicated time points, as measured by MTT cell viability assay (n=5 per group). Two-way ANOVA; p=0.803 (4T1), p=0.124 (EO771), p=0.446 (PyMT). **i**, Apoptosis of control (sgNTC) or Cpt1a-deficient (sgCpt1a) 4T1 or EO771 cells was assessed by Annexin V and 7-AAD staining (n=3). Representative dot plots of 4T1 cells are shown, and unpaired Student's t-tests were performed. **j**, Experimental timeline for *in vivo* lung metastasis evaluation of control (sgNTC) and Cpt1a-*KO* (sgCpt1a) 4T1 or EO771 tumor cells. Primary tumors were resected on day 14-18 post-implantation, and whole lungs were collected 4 weeks and 8 weeks after 4T1 or EO771 tumor resection, respectively. **k**, Primary tumor growth from mice described in (**j**) (n≥5 mice per group). Two-way ANOVA; p=0.369 (4T1), p=0.727 (EO771). **l-m**, Lung metastasis of mice described in (**j**). Number of lung surface metastases in BALB/c-Cas9 mice or BL6-Cas9 (**l**) were determined. Representative H&E images are shown in (**m**). Welch's t-tests were performed. **k**, The D454 site is conserved in human and mouse CPT1A, located within the catalytic carnitine O-palmitoyltransferase domain. The D454G mutant in humans leads to low CPT1A activity. **o**, Immunoblot of CPT1A in Ctrl (sgNTC) (lane 1) or *Cpt1a*-KO (sgCpt1a) cells transduced with empty vector (EV) (lane 2), WT CPT1A (lane 3), or CPT1A D454G (lane 4). **p**, Basal respiration OCR was determined in sgNTC control (Ctrl), *Cpt1a*-KO (KO) 4T1 cells or KO cells re-expressing WT or D454G CPT1A, per $5 \times 10^4$ cells from Figure 1g. One-way ANOVA ($p=8.19 \times 10^{-7}$) with Tukey's post hoc. *p<0.05, **p<0.01, ***p<0.005, ****p<0.001. ns, not significant.

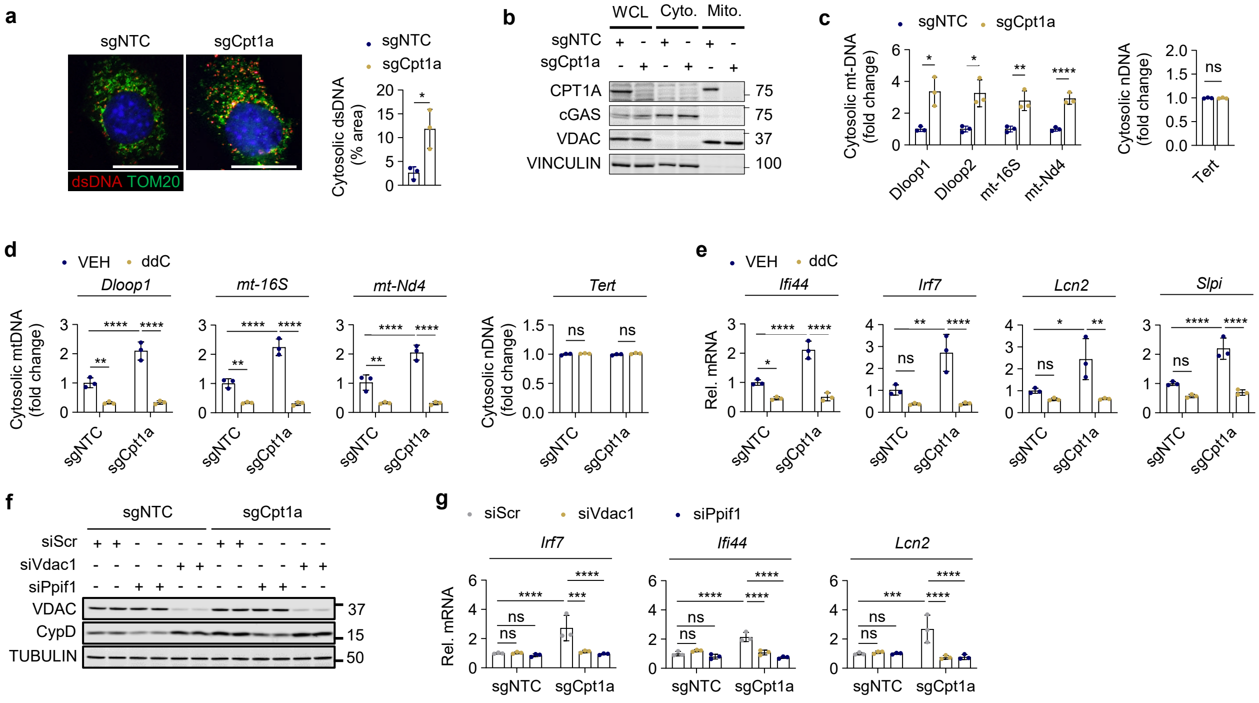

**Extended Data.3 | The mPTP pore is required for CPT1A loss driven inflammation. a**, Immunofluorescence staining of dsDNA (red) and Tom20 (green) in control (sgNTC) and *Cpt1a*-KO (sgCpt1a) PyMT cells. Scale bar 20μM. Cytosolic dsDNA coverage was determined as a percent (%) of total area. Unpaired Student's t-test was performed. **b-c**, Fractionation of PyMT control (Ctrl sg or sgNTC) or *Cpt1a*-KO (Cpt1a sg or sgCpt1a) cells was performed to collect the cytosolic fraction. **b**, Immunoblot to demonstration fractionation. Markers to denote mitochondrial (Mito., CPT1A and VDAC) and cytosolic (Cyto., cGAS and VINCULIN) fractions are shown. Whole cell lysates (WCL) serve as an input control. **c**, Relative levels of mitochondrial DNA (mtDNA) regions (Dloop1, Dloop2, mt-16S, mt-Nd4) (left) or nuclear DNA (nDNA, Tert) (right) in the cytosolic fraction was determined using RT-PCR. Unpaired Student's t-tests were performed. **d-e**, Control (sgNTC) or *Cpt1a*-KO (sgCpt1a) PyMT cells were treated with vehicle (VEH) or dideoxycytidine (ddC) to deplete mtDNA. n=3 per group. **d**, Relative levels of mtDNA regions (left) or nDNA (right) in the cytosolic fraction was determined using RT-PCR. Two-way ANOVA with Tukey's post hoc; p=$8.48\times10^{-4}$ (Dloop1), p=$1.70\times10^{-4}$ (mt-16S), p=0.00129 (mt-Nd4), p=0.757 (Tert). **e**, Relative expression of *Ifi44, Irf7, Lcn2*, and *Slpi* was determined by RT-PCR. Two-way ANOVA with Tukey's post hoc; p=$7.57\times10^{-4}$ (Ifi44), p=0.0103 (Irf7), p=0.0319 (Lcn2), p=0.00128 (Slpi). **f-g**, Control (sgNTC) or *Cpt1a*-KO (sgCpt1a) 4T1 cells were transfected with scrambled control (Ctrl), Vdac1, or CypD (Ppif) si RNA. **f**, Immunoblot confirms CypD and VDAC levels. Tubulin serves as loading control. **g**, Relative expression of *Irf7, Ifi44,* and *Lcn2* was determined by RT-PCR. Two-way ANOVA with Tukey's post hoc; p=0.00231 (Irf7), p=$6.37\times10^{-5}$ (Ifi44), p=0.00132 (Lcn2 *p<0.05, **p<0.01, ***p<0.005, ****p<0.001. ns, not significant.

**Supplementary Table 1. Composition of the metabolism-focused sgRNA library and single gene KO sgRNA used in this study**

| Gene Symbol | sgRNA ID | sgRNA Sequence (5'→3') |
|---|---|---|
| Cpt1a | Cpt1a_sg1 | CACATTGTCGTGTACCACAG |
| Cpt1a | Cpt1a_sg2 | CATACTGCTGTATCGTCGCA |
| Cpt1a | Cpt1a_sg3 | ACCTTGGACCCAAATTGCAG |
| Cpt1a | Cpt1a_sg4 | ACGTTGGACGAATCGGAACA |
| Ppif | Ppif_sg1 | GATGTCGTGCCAAAGACTGC |
| Ppif | Ppif_sg2 | CCGCTCGTGTACTTGGACGT |
| NTC | NTC_sg1 | AAAAAGTCCGCGATTACGTC |
| NTC | NTC_sg2 | AAAACGGCTCGATCGGTGAT |
| NTC | NTC_sg3 | AAAACGTAATTATACCGAGC |
| NTC | NTC_sg4 | AAAATTGCACCTTCCCGGCC |

**Supplementary Table 2. Composition of the metabolism-focused sgRNA library and single gene KO sgRNA used in this study**

| Gene Symbol | sgRNA ID | sgRNA Sequence (5'→3') |
|---|---|---|
| Acaa1b | Acaa1b_sg1 | AACCACTGTCCTGAATGACA |
| Acaa1b | Acaa1b_sg2 | GGCGGGAAGAAATATTCCCA |
| Acaa1b | Acaa1b_sg3 | TGTGTGCACTAGGATAACTT |
| Acaa1b | Acaa1b_sg4 | GGAGCCAGAGAGATTACCTC |
| Acaa2 | Acaa2_sg1 | GCCCACGATGACACTATCGA |
| Acaa2 | Acaa2_sg2 | GCTGAGGTCGTCTTGTGTGG |
| Acaa2 | Acaa2_sg3 | CGAGGCTGGCTACTTCAATG |
| Acaa2 | Acaa2_sg4 | CGTCCGTGTTCAAGAAAGAC |
| Acaca | Acaca_sg1 | GCCATTCATTATCACTACGT |
| Acaca | Acaca_sg2 | AATGCATGCGATCTATCCGT |
| Acaca | Acaca_sg3 | TTGATTCATAGGTACCGAAG |
| Acaca | Acaca_sg4 | AAGCCCTTCGAACATACACC |
| Acly | Acly_sg1 | GAGAGAGATTGACCCCGACG |
| Acly | Acly_sg2 | AGAGCGATTCGAGATTACCA |
| Acly | Acly_sg3 | TTGTCACCTGTACACGACGG |
| Acly | Acly_sg4 | GGACGAAAAGCTGAATACCG |
| Aco1 | Aco1_sg1 | CCACCGCCATTTAGGCCGCG |
| Aco1 | Aco1_sg2 | CATATGCTATTACCAGAGGG |
| Aco1 | Aco1_sg3 | TAGCCCACCACATCAAACCT |
| Aco1 | Aco1_sg4 | GTGTAGCACCTCCGACAAGT |
| Acss2 | Acss2_sg1 | GCTGGGAACCTACTACCCGG |
| Acss2 | Acss2_sg2 | CAGAACGCCGGTGCAGCTCG |
| Acss2 | Acss2_sg3 | AAGGGAAAATATTCACTGAG |
| Acss2 | Acss2_sg4 | GCATTGTGGTCAAACATCTG |
| Adpgk | Adpgk_sg1 | CTCTCACGACCTCTCCAACG |
| Adpgk | Adpgk_sg2 | ATGCTGCTTTAATTGGACAG |
| Adpgk | Adpgk_sg3 | CCAGGTACTCTAAAATAAGG |
| Adpgk | Adpgk_sg4 | GCTGGCCAGTATGACCAACA |
| Ahcy | Ahcy_sg1 | GCGCACCTGACAGAAGCTGT |
| Ahcy | Ahcy_sg2 | TGTCAACGATTCTGTCACCA |
| Ahcy | Ahcy_sg3 | TGACCCTATCATACCCTCCA |
| Ahcy | Ahcy_sg4 | TGTGATGATTGCGGGCAAGG |
| Ahr | Ahr_sg1 | AGCTGTGCACAAGAGGATCG |
| Ahr | Ahr_sg2 | GTATAATAGACTGCTGCCTG |
| Ahr | Ahr_sg3 | TCTCCGGTAGCAAACATGAA |
| Ahr | Ahr_sg4 | GTGGAAAGAATCCTTACTTG |

| | | |
|---|---|---|
| Aldob | Aldob_sg1 | TATCCACAGTTGGACCAAGG |
| Aldob | Aldob_sg2 | AATTCCATTAGCCAGAGCAT |
| Aldob | Aldob_sg3 | CCGCCTGCAAAGGATAAAGG |
| Aldob | Aldob_sg4 | GGTCCCTATTGTTGAGCCAG |
| Arg1 | Arg1_sg1 | AATAAACTTACTGTTCCCCA |
| Arg1 | Arg1_sg2 | AGTATGACGTGAGAGACCAC |
| Arg1 | Arg1_sg3 | AAATGACACATAGGTCAGGG |
| Arg1 | Arg1_sg4 | AGATGTACCAGGATTCTCCT |
| Arg2 | Arg2_sg1 | CCAATGTACACAATATTTGG |
| Arg2 | Arg2_sg2 | TACACAACCAGATTATTGTA |
| Arg2 | Arg2_sg3 | TTTCCAGATACAGTGGTGAG |
| Arg2 | Arg2_sg4 | GTCCACTCTGTAGCTATAGT |
| Ass1 | Ass1_sg1 | TAACATCTTACCTTAATCTG |
| Ass1 | Ass1_sg2 | TGGGATCCTGGAAAACCCCA |
| Ass1 | Ass1_sg3 | AAGGCACTTCCTCACCAGGT |
| Ass1 | Ass1_sg4 | TGCCTTCACCTGTAGCAACA |
| Atg16l1 | Atg16l1_sg1 | CATACTTACGAAGACATACG |
| Atg16l1 | Atg16l1_sg2 | CGAACTGCACAAGAAGCGTG |
| Atg16l1 | Atg16l1_sg3 | GAAACTGAGGAAAACTACTG |
| Atg16l1 | Atg16l1_sg4 | TGCAAGCCGAATCTGGACTG |
| Atg3 | Atg3_sg1 | GTAGATACATATCACAACAC |
| Atg3 | Atg3_sg2 | GACAGGCTACCCTAGACACA |
| Atg3 | Atg3_sg3 | GGGTGTAATCACCCCAGAAG |
| Atg3 | Atg3_sg4 | TAACAGTTCCATGCTACAAG |
| Atg5 | Atg5_sg1 | AAGAGTCAGCTATTTGACGT |
| Atg5 | Atg5_sg2 | AAATGTACTGTGATGTTCCA |
| Atg5 | Atg5_sg3 | CCTTCTACACTGTCCATCCA |
| Atg5 | Atg5_sg4 | AAGAAAAACTCACCATTTCA |
| Atg7 | Atg7_sg1 | TCTCCTACTCCAATCCCGTG |
| Atg7 | Atg7_sg2 | TGGGGTCCATACATCCACTG |
| Atg7 | Atg7_sg3 | CTTAAAAGCCTCAAGTGTGT |
| Atg7 | Atg7_sg4 | CTTGAATAAGAAGTAGGGCA |
| Atp5a1 | Atp5a1_sg1 | TGGTCAGAAGCGGTCCACTG |
| Atp5a1 | Atp5a1_sg2 | GCTCCCGCACAGAGATTCGG |
| Atp5a1 | Atp5a1_sg3 | ACTGGGCGTGTGTTAAGCAT |
| Atp5a1 | Atp5a1_sg4 | CCAACAGCTCCTCGCCAACG |
| Atp5b | Atp5b_sg1 | CTGCTGGCCCCATACGCCAA |
| Atp5b | Atp5b_sg2 | GCGCTTACCAGGATGAACCC |
| Atp5b | Atp5b_sg3 | AAATACAGAGTAACCACCAT |
| Atp5b | Atp5b_sg4 | CCCACCCTAGCCACCGACAT |

| | | |
|---|---|---|
| Atp6ap1 | Atp6ap1_sg1 | GAGGATTTCACAGCATACGG |
| Atp6ap1 | Atp6ap1_sg2 | GATATGACCCTCATGTGTGT |
| Atp6ap1 | Atp6ap1_sg3 | TAGCTAGATCCACATGCAAG |
| Atp6ap1 | Atp6ap1_sg4 | GTGTCATTGTAACTCACAGG |
| Batf | Batf_sg1 | AGAGATCAAACAGCTCACCG |
| Batf | Batf_sg2 | AGGACTCATCTGATGATGTG |
| Batf | Batf_sg3 | GTGGGTACTCACCAGGTGAA |
| Batf | Batf_sg4 | AGGGGGTACCTGTTTGCCAG |
| Bcl6 | Bcl6_sg1 | TCTCCACGACCTCACGACCT |
| Bcl6 | Bcl6_sg2 | ATGTTGCTGTGACACAAACT |
| Bcl6 | Bcl6_sg3 | AATGCACCCTTGAACCGGAA |
| Bcl6 | Bcl6_sg4 | GAGGGAAGGCAATATCATGG |
| Cd5l | Cd5l_sg1 | GGCTGATATGATGCGCCACG |
| Cd5l | Cd5l_sg2 | CAACGGAACGGAAGACACGT |
| Cd5l | Cd5l_sg3 | AGCTGCAACAAGAATACTCA |
| Cd5l | Cd5l_sg4 | ACCAAAGTGCAGCTAGTGGG |
| Cers4 | Cers4_sg1 | TTGAGAATCTTACAACCCTG |
| Cers4 | Cers4_sg2 | CGCTTCGGCAGACTCAACGC |
| Cers4 | Cers4_sg3 | GTTACCACCCAATGTCACAT |
| Cers4 | Cers4_sg4 | GCGGAGCAAGTTGACGCTGT |
| Cox10 | Cox10_sg1 | TCTGTCCCGGAAGCCAAATG |
| Cox10 | Cox10_sg2 | GGGAGTGAATCCACTCACAG |
| Cox10 | Cox10_sg3 | TATACAGGGATTGCCACACA |
| Cox10 | Cox10_sg4 | AGTTGGCAGCACAGGATGCG |
| Cox15 | Cox15_sg1 | AGAAAGGGTTGGCTCAACCG |
| Cox15 | Cox15_sg2 | AGGCGGTACTGACTGACCCG |
| Cox15 | Cox15_sg3 | GGTACATGGAATACTCACAC |
| Cox15 | Cox15_sg4 | TAGGGTGGCGTCCAGAACAC |
| Cox7a1 | Cox7a1_sg1 | GACCTCCCAGTACACTTGAA |
| Cox7a1 | Cox7a1_sg2 | AAGCCACTTAGAAAACCGTG |
| Cox7a1 | Cox7a1_sg3 | TGGTAGATGAGCTAAAAGAC |
| Cox7a1 | Cox7a1_sg4 | AAAAGACCGGACCAGAGCCT |
| Cps1 | Cps1_sg1 | TGAGCCTCACAATTTCGTCG |
| Cps1 | Cps1_sg2 | ATGCAGACCGAATCATCACA |
| Cps1 | Cps1_sg3 | TACAGTATTCCATGGAAGTG |
| Cps1 | Cps1_sg4 | GTTGGTGGCATCTCGTGTCG |
| Cpt1a | Cpt1a_sg1 | CACATTGTCGTGTACCACAG |
| Cpt1a | Cpt1a_sg2 | CATACTGCTGTATCGTCGCA |
| Cpt1a | Cpt1a_sg3 | ACCTTGGACCCAAATTGCAG |
| Cpt1a | Cpt1a_sg4 | ACGTTGGACGAATCGGAACA |

| | | |
|---|---|---|
| Dgat2 | Dgat2_sg1 | GATCTGCCCTGTCACGCGAG |
| Dgat2 | Dgat2_sg2 | CTGGCTCAACAGATCTAAGG |
| Dgat2 | Dgat2_sg3 | AAGGCCCTATTTGGCTACGT |
| Dgat2 | Dgat2_sg4 | GTCTCGGAAGTAGCGCCACA |
| Dhfr | Dhfr_sg1 | GACATGGTTTGGATAGTCGG |
| Dhfr | Dhfr_sg2 | AACCTCAGAGAACCACCACG |
| Dhfr | Dhfr_sg3 | TCGCCGTGTCCCAAAATATG |
| Dhfr | Dhfr_sg4 | CAGCCCGGCCAATACCTGAG |
| Fasl | Fasl_sg1 | AGGACCACAACACAAATCTG |
| Fasl | Fasl_sg2 | CTTCACTCCAGAGATCAGAG |
| Fasl | Fasl_sg3 | CCTCTGAAAAAAAAGAGCCG |
| Fasl | Fasl_sg4 | GGAACTGGCAGAACTCCGTG |
| Fasn | Fasn_sg1 | CTACCAGGCCATCCGTAGTG |
| Fasn | Fasn_sg2 | TGTCTCCGAAAAGAGCCGGG |
| Fasn | Fasn_sg3 | TTGGTGGAGCCAATTAACAG |
| Fasn | Fasn_sg4 | ACTGGCAATCTGATTGTGAG |
| Fbp1 | Fbp1_sg1 | CATGGCAAGGACCAACATGG |
| Fbp1 | Fbp1_sg2 | CACAAGAACACAGGTAGCGT |
| Fbp1 | Fbp1_sg3 | TGGCTCAACCAATGTGACTG |
| Fbp1 | Fbp1_sg4 | AACATCTACAGCCTTAATGA |
| Foxp3 | Foxp3_sg1 | CATACCTGATGCATGAAGTG |
| Foxp3 | Foxp3_sg2 | TCTACCCACAGGGATCAATG |
| Foxp3 | Foxp3_sg3 | AGGTCGGGACCTGCGAAGTG |
| Foxp3 | Foxp3_sg4 | GCAAGAGCTCTTGTCCATTG |
| G6pdx | G6pdx_sg1 | AGAGGTGGAAACTGACAACG |
| G6pdx | G6pdx_sg2 | TGCCCGCTCACGACTCACAG |
| G6pdx | G6pdx_sg3 | ATGACCCCACAGTACCCCAT |
| G6pdx | G6pdx_sg4 | AGAGATGGTCCAGAATCTCA |
| Gapdh | Gapdh_sg1 | GCTGTGGCGTGATGGCCGTG |
| Gapdh | Gapdh_sg2 | AAACAGGCCCACTTGAAGGG |
| Gapdh | Gapdh_sg3 | TGCCATTTGCAGTGGCAAAG |
| Gapdh | Gapdh_sg4 | GGCCGGTGCTGAGTATGTCG |
| Gata3 | Gata3_sg1 | CTACTACGGAAACTCCGTCA |
| Gata3 | Gata3_sg2 | CCGGGTTCGGATGTAAGTCG |
| Gata3 | Gata3_sg3 | GCAGCTGCACCTGATACTTG |
| Gata3 | Gata3_sg4 | TCCAAGACGTCCATCCACCA |
| Gclc | Gclc_sg1 | TGTGCCGGTCCTTGACTGCG |
| Gclc | Gclc_sg2 | CAATATGAGGAAACGCCGGA |
| Gclc | Gclc_sg3 | AGAAACATCCGGCATCGGAG |
| Gclc | Gclc_sg4 | TGTAGATGATAGAACACGGG |

| | | |
|---|---|---|
| Glo1 | Glo1_sg1 | CGATCCAGACCCTAGCACCA |
| Glo1 | Glo1_sg2 | GCACTGCGTGAGCTCAAGGG |
| Glo1 | Glo1_sg3 | GGATAAGAACGATATCCCCA |
| Glo1 | Glo1_sg4 | GAGACTCAGAGTTACCACAA |
| Gls | Gls_sg1 | CGACGCGTTCGGCAACAGCG |
| Gls | Gls_sg2 | TGTACATCGCTATGTTGGGA |
| Gls | Gls_sg3 | GATTGCGAACATCTGATCCC |
| Gls | Gls_sg4 | ATATAACTCATCGATGTGTG |
| Gls2 | Gls2_sg1 | CGTCCGGTACTACCTCGGTG |
| Gls2 | Gls2_sg2 | GGGGATCGGAATTACGCCAT |
| Gls2 | Gls2_sg3 | AAAAGCAGGTCACCAAGTCG |
| Gls2 | Gls2_sg4 | TGAGTCAGGCAGTGTCATGG |
| Got1 | Got1_sg1 | GATCCCCGCAAGGTTAACCT |
| Got1 | Got1_sg2 | GTTGGTGATGATACGTAGAT |
| Got1 | Got1_sg3 | AGACCTAGAGAAAGATGCGT |
| Got1 | Got1_sg4 | CATTCGGCCCTATTGCTACT |
| Got2 | Got2_sg1 | TGGAGGTCCCATTTCAACAT |
| Got2 | Got2_sg2 | TTTCTGCCCAAACCATCCTG |
| Got2 | Got2_sg3 | CATCCTCCTCACCTTCACCA |
| Got2 | Got2_sg4 | AGCTCACCTTCCGGACACTG |
| Gpt2 | Gpt2_sg1 | GCGGTGGAGTACGCTGTGCG |
| Gpt2 | Gpt2_sg2 | ACGCTAAGAAACGAGCGCGG |
| Gpt2 | Gpt2_sg3 | GTTCTCTGCATTATCAACCC |
| Gpt2 | Gpt2_sg4 | GGGGATGGGAATCATCACGC |
| Hif1a | Hif1a_sg1 | TGAACATCAAGTCAGCAACG |
| Hif1a | Hif1a_sg2 | ATAACGTGAACAAATACATG |
| Hif1a | Hif1a_sg3 | GTGAGAAAACTTCTGGATGC |
| Hif1a | Hif1a_sg4 | AGTAAGAAAATTTCATATCG |
| Hk1 | Hk1_sg1 | CCGACAATCCAAAATAGACG |
| Hk1 | Hk1_sg2 | CGTAGCCGCCATTGAAACGT |
| Hk1 | Hk1_sg3 | GGATCTTTACCAGTAGGACT |
| Hk1 | Hk1_sg4 | CTCCCGGGATTATAACCCAA |
| Hk2 | Hk2_sg1 | ATTCCCGAGGACATCATGCG |
| Hk2 | Hk2_sg2 | GGAGATGCGTCACATTGACA |
| Hk2 | Hk2_sg3 | ATCCGGAGTTGACCTCACAA |
| Hk2 | Hk2_sg4 | GGAGTGGCACACACATAAGT |
| Idh1 | Idh1_sg1 | CCCAGCCTGTCACTAGCCGG |
| Idh1 | Idh1_sg2 | GGCTATAAAGAAATACAACG |
| Idh1 | Idh1_sg3 | AATTCAAGTTGAAACAAATG |
| Idh1 | Idh1_sg4 | TGGTACATGACTTTGAAGGT |

| | | |
|---|---|---|
| Idh2 | Idh2_sg1 | GGCCACCCAGAAGTACAGTG |
| Idh2 | Idh2_sg2 | TCGAGCTGGCACGTTCAAGT |
| Idh2 | Idh2_sg3 | TCACCGTCCATCTCCACTAC |
| Idh2 | Idh2_sg4 | ACATCGGCTCATCGACGACA |
| Ikzf2 | Ikzf2_sg1 | CCTAATTGAGAGCAGCGAGG |
| Ikzf2 | Ikzf2_sg2 | GCTTGTCATGTGACTTGGCG |
| Ikzf2 | Ikzf2_sg3 | TATGAACTTAACATATGAGA |
| Ikzf2 | Ikzf2_sg4 | GGGTAAAAGAAGCTCCGCAC |
| Il2ra | Il2ra_sg1 | GTGTCTGTATGACCCACCCG |
| Il2ra | Il2ra_sg2 | ATCTTGCAGATGCTAATAGC |
| Il2ra | Il2ra_sg3 | GAGAGGTTTCCGAAGACTAA |
| Il2ra | Il2ra_sg4 | GAATCTTCATGTTTCCAAGG |
| Kdsr | Kdsr_sg1 | ACCATGAAGGAGCGACGGGT |
| Kdsr | Kdsr_sg2 | ACAGTGACGTACACATTGTA |
| Kdsr | Kdsr_sg3 | AGTGGAGAATGTCATAAAGC |
| Kdsr | Kdsr_sg4 | GCTATTGAGTGCTACAAACA |
| Ldha | Ldha_sg1 | CAAGCTGGTCATTATCACCG |
| Ldha | Ldha_sg2 | GTTGCAATCTGGATTCAGCG |
| Ldha | Ldha_sg3 | GGAGAACATGGCGACTCCAG |
| Ldha | Ldha_sg4 | GTCATGGAAGACAAACTCAA |
| Mdh1 | Mdh1_sg1 | GTCAGCGCCATCGATCCCCA |
| Mdh1 | Mdh1_sg2 | GTCCATAGATGTCATTGCAA |
| Mdh1 | Mdh1_sg3 | GACATTCTTTACATCATCAG |
| Mdh1 | Mdh1_sg4 | GTCTTTGGGAAAGACCAGGT |
| Mthfd1 | Mthfd1_sg1 | ACACCAACGATAGATTCCTG |
| Mthfd1 | Mthfd1_sg2 | CACTATGAATCCGTGCACAG |
| Mthfd1 | Mthfd1_sg3 | GATTGCCGGAAGGCACGCGG |
| Mthfd1 | Mthfd1_sg4 | GGTAGCGTCCAGTAAGAAAG |
| Mthfd2 | Mthfd2_sg1 | TCGATGAGATATTGTGACTG |
| Mthfd2 | Mthfd2_sg2 | AGATAATTAAGCGAACAGGT |
| Mthfd2 | Mthfd2_sg3 | GCTTTCATGTCATTAACGTG |
| Mthfd2 | Mthfd2_sg4 | CTATGTTCTCAACAAAACCA |
| Ndufc1 | Ndufc1_sg1 | TTGGCAGTTGGACTGTCCGT |
| Ndufc1 | Ndufc1_sg2 | CGAAAACGAGCGCAGCACTA |
| Ndufc1 | Ndufc1_sg3 | CACGGTCGAAGTTCTATGTC |
| Ndufc1 | Ndufc1_sg4 | CAATGCCAAACCTAACTGGT |
| Nfe2l2 | Nfe2l2_sg1 | TGAAGACTGAACTTTCAGCG |
| Nfe2l2 | Nfe2l2_sg2 | GTTCTGTTTGACACTTCCAG |
| Nfe2l2 | Nfe2l2_sg3 | TTCAACCCGAAGCACGCTGA |
| Nfe2l2 | Nfe2l2_sg4 | GGTGGGATTTGAGTCTAAGG |

| | | |
|---|---|---|
| Olah | Olah_sg1 | AAAACCAGAACTTACGTGAG |
| Olah | Olah_sg2 | TGCATGCTGTAAGACTGGCT |
| Olah | Olah_sg3 | TTACAAGATCTAAATACCTG |
| Olah | Olah_sg4 | ATTAATCTTTCGGCCCCACT |
| Otc | Otc_sg1 | CAGTCCATTGACAATTGGGA |
| Otc | Otc_sg2 | CCTTCAAGCAGCTACTCCAA |
| Otc | Otc_sg3 | TAGAAAGGGTCACACTTCTG |
| Otc | Otc_sg4 | AAATTCAGGATCAAGCAGAA |
| Pck1 | Pck1_sg1 | ACTGACAGACTCGCCCTATG |
| Pck1 | Pck1_sg2 | GTGGCCGAGACTAGCGATGG |
| Pck1 | Pck1_sg3 | CCTTTGGAAGCGGATATGGT |
| Pck1 | Pck1_sg4 | TCGCAGATGTGGATATACTC |
| Pck2 | Pck2_sg1 | TGCGTATTATGACCCGCCTG |
| Pck2 | Pck2_sg2 | TGATTGTAACTCCTTCGCAG |
| Pck2 | Pck2_sg3 | AGGGTTTGGATGCTACGGCA |
| Pck2 | Pck2_sg4 | ATGGAAGCACATACATAATG |
| Pdcd1 | Pdcd1_sg1 | CAATACAGGGATACCCACTA |
| Pdcd1 | Pdcd1_sg2 | GACACACGGCGCAATGACAG |
| Pdcd1 | Pdcd1_sg3 | CAGCTTGTCCAACTGGTCGG |
| Pdcd1 | Pdcd1_sg4 | GCTCAAACCATTACAGAAGG |
| Pdk1 | Pdk1_sg1 | TTGTCGCAGAAACATAAACG |
| Pdk1 | Pdk1_sg2 | ATGGCTATGAGAACGCTAGG |
| Pdk1 | Pdk1_sg3 | TTGATAGCCTTATTGTTCGG |
| Pdk1 | Pdk1_sg4 | AAACACCATGTGATAGAGAT |
| Pfkm | Pfkm_sg1 | CCTCACGGTAGAGCGAACAG |
| Pfkm | Pfkm_sg2 | GCGCCTTGGATATGACACCC |
| Pfkm | Pfkm_sg3 | TTAGACCAAAGACGTGACCA |
| Pfkm | Pfkm_sg4 | CATAGACACGCTCTCCCACG |
| Pgk1 | Pgk1_sg1 | TAAGGTGCTCAACAACATGG |
| Pgk1 | Pgk1_sg2 | TCAAGAACAGAACATCCCTG |
| Pgk1 | Pgk1_sg3 | GGACTGCACACCGAGCCCAT |
| Pgk1 | Pgk1_sg4 | CTTCCTCTACATGAAAGCGG |
| Pgk2 | Pgk2_sg1 | GGATACCATCAGGCCGACCG |
| Pgk2 | Pgk2_sg2 | GGATGATAGACCCATTATCT |
| Pgk2 | Pgk2_sg3 | CGGGCTCACAGTTCTACGGT |
| Pgk2 | Pgk2_sg4 | GGAAGGCTTCTACTTTAGCA |
| Pgm1 | Pgm1_sg1 | CATTACCGATGGACGCGCTG |
| Pgm1 | Pgm1_sg2 | ATCATCTCTCCCCACGATCG |
| Pgm1 | Pgm1_sg3 | TGGGGGTTATATCAGAGAAG |
| Pgm1 | Pgm1_sg4 | AGGCCAACTGCACAAACTCG |

| | | |
|---|---|---|
| Pgm2 | Pgm2_sg1 | CGGCCGCTTCTACATGACCG |
| Pgm2 | Pgm2_sg2 | CGCATAGACGCCATGCACGG |
| Pgm2 | Pgm2_sg3 | CAGCCAGCCATAATCCAGGA |
| Pgm2 | Pgm2_sg4 | CAGCAGCATAGGTGAGATTG |
| Pik3c3 | Pik3c3_sg1 | AGCCTGTAAGAACTCAACAC |
| Pik3c3 | Pik3c3_sg2 | ATACACATCCCATATAGTCA |
| Pik3c3 | Pik3c3_sg3 | CTCACCAAGGCTCATCGGCA |
| Pik3c3 | Pik3c3_sg4 | ATGGACCAGGCGATCTACAA |
| Pkm | Pkm_sg1 | TTTCTCTCATGGAACCCATG |
| Pkm | Pkm_sg2 | TGAAATAGCACATGCCTGTG |
| Pkm | Pkm_sg3 | GGGCAGAGTCAATGTCCAGG |
| Pkm | Pkm_sg4 | CTTCCTGACTTCATGCACGT |
| Ppat | Ppat_sg1 | ACCTTGGAATCGGACATACG |
| Ppat | Ppat_sg2 | ATAAGACGCCCGATGCAGAG |
| Ppat | Ppat_sg3 | TGATCACTCTGGGACTCGTG |
| Ppat | Ppat_sg4 | AGGGGTGTATGCGAGTAACT |
| Prdx2 | Prdx2_sg1 | ATCAAGCTTTCGGACTACAG |
| Prdx2 | Prdx2_sg2 | CCTTCAGGATCAATACCCCA |
| Prdx2 | Prdx2_sg3 | GCTAAAAGCGATGATCTCCG |
| Prdx2 | Prdx2_sg4 | CTTCCGAAAGCTAGGCTGCG |
| Rheb | Rheb_sg1 | AACAAACTGAATTGTCAATG |
| Rheb | Rheb_sg2 | CCATATCCAACAACTTGCCA |
| Rheb | Rheb_sg3 | TTCAGCTTGTAGACACAGCG |
| Rheb | Rheb_sg4 | TCATAGGATACCTATTATGT |
| Rorc | Rorc_sg1 | CTTGAGTATAGTCCAGAACG |
| Rorc | Rorc_sg2 | GTCATCTGGGATCCACTACG |
| Rorc | Rorc_sg3 | TCTGGGGCACTGCAGAAACT |
| Rorc | Rorc_sg4 | GACAAGCAGAGGCCTCGGGT |
| Sdha | Sdha_sg1 | GTCAGTTACCTCAACCACAG |
| Sdha | Sdha_sg2 | TTCTACTCAATACCCAGTGG |
| Sdha | Sdha_sg3 | TGCACAGTGCAATGACACCA |
| Sdha | Sdha_sg4 | ACTGTGCATTACAACATGGG |
| Sdhb | Sdhb_sg1 | TGCGCCATGAACATCAACGG |
| Sdhb | Sdhb_sg2 | ACAGTATCTGCAGTCCATCG |
| Sdhb | Sdhb_sg3 | ACCTCGAATGCAGACGTACG |
| Sdhb | Sdhb_sg4 | TAGAAGTTACTCAAATCCTG |
| Sgpp1 | Sgpp1_sg1 | TGCCTAAGTAGAATCTACAT |
| Sgpp1 | Sgpp1_sg2 | TGAGCAGGAACATGGCGATG |
| Sgpp1 | Sgpp1_sg3 | CCTCGCCCGTCAACGAGTTG |
| Sgpp1 | Sgpp1_sg4 | TGGGTGCTGGTCATGTACCT |

| | | |
|---|---|---|
| Shmt1 | Shmt1_sg1 | TGTAGAATATCATACCAGCA |
| Shmt1 | Shmt1_sg2 | TCGGCTGGCAAAATTCTCCG |
| Shmt1 | Shmt1_sg3 | CCCGGAACCTGGACTACGCA |
| Shmt1 | Shmt1_sg4 | AAGCCCATGATTCGCCCATG |
| Shmt2 | Shmt2_sg1 | CGGCAGATACTACGGAGGAG |
| Shmt2 | Shmt2_sg2 | AACATCCGCGTACTTGAAAG |
| Shmt2 | Shmt2_sg3 | AGCCTCATGATCGAATCATG |
| Shmt2 | Shmt2_sg4 | TAGTCGATGAGGCCAGTTTG |
| Slc16a1 | Slc16a1_sg1 | ACTACTAAGAAAGACCAAAG |
| Slc16a1 | Slc16a1_sg2 | CACCAGCGATCATTACTGGA |
| Slc16a1 | Slc16a1_sg3 | GACTTGCAGCCAACACCAAG |
| Slc16a1 | Slc16a1_sg4 | AGGCCCTATTGGTCTCATCA |
| Slc16a7 | Slc16a7_sg1 | ATTACCTCCAATGAAGCCAA |
| Slc16a7 | Slc16a7_sg2 | AGAGGTACTGGATTCGTGGA |
| Slc16a7 | Slc16a7_sg3 | GCTCAGTACGCTAAACACAT |
| Slc16a7 | Slc16a7_sg4 | TTCACCAACACACTACTGAT |
| Slc1a5 | Slc1a5_sg1 | AATCCCTATCGATTCCTGTG |
| Slc1a5 | Slc1a5_sg2 | TACAACAGAGTCGTTGATGG |
| Slc1a5 | Slc1a5_sg3 | GCGGGAGATCAATTCAACCA |
| Slc1a5 | Slc1a5_sg4 | GTGGTGTGCAGCCTGATCGG |
| Slc20a1 | Slc20a1_sg1 | GTAGAAAGGTTACCTTACGG |
| Slc20a1 | Slc20a1_sg2 | TCAGTATCACACCGTGCACA |
| Slc20a1 | Slc20a1_sg3 | CCGGAACGGCTTGATAGATG |
| Slc20a1 | Slc20a1_sg4 | GCCACATATTGCCATAGTGT |
| Slc2a1 | Slc2a1_sg1 | CCTGCTCATCAATCGTAACG |
| Slc2a1 | Slc2a1_sg2 | TCAGCATGGAGTTCCGCCTG |
| Slc2a1 | Slc2a1_sg3 | GTGTCACCTACAGCTCTACG |
| Slc2a1 | Slc2a1_sg4 | CAAACATGGAACCACCGCTA |
| Slc38a1 | Slc38a1_sg1 | ATACTTTGGTGTGCACGCGT |
| Slc38a1 | Slc38a1_sg2 | TGCATGGTGTATGAGAAGCT |
| Slc38a1 | Slc38a1_sg3 | TCACCATCACCACCAACACT |
| Slc38a1 | Slc38a1_sg4 | AGATTGGCAGGACGGACGGG |
| Slc38a2 | Slc38a2_sg1 | CCACCAAAGCAGCTTCCACG |
| Slc38a2 | Slc38a2_sg2 | CTCAAGACTGCCAACGAAGG |
| Slc38a2 | Slc38a2_sg3 | GCAGTGACAATGGAAGAATG |
| Slc38a2 | Slc38a2_sg4 | GAGTTGAAGATGAAATAGCG |
| Slc3a2 | Slc3a2_sg1 | GTTCACCGGCTTATCCAAGG |
| Slc3a2 | Slc3a2_sg2 | CGCCCGAACGATGATAACCA |
| Slc3a2 | Slc3a2_sg3 | TATCACCAAGAACTTAAGTG |
| Slc3a2 | Slc3a2_sg4 | GTACTGAATCCCTAGTCACT |

| | | |
|---|---|---|
| Slc6a1 | Slc6a1_sg1 | CACCAACATGACCAGCGCCG |
| Slc6a1 | Slc6a1_sg2 | GCAGAAATACACGAGCACCC |
| Slc6a1 | Slc6a1_sg3 | TACCTCTGTGGGAAAAACGG |
| Slc6a1 | Slc6a1_sg4 | TCCATGTGTCCCGGTCAGGG |
| Slc7a1 | Slc7a1_sg1 | GCCATGGCATAGATAACTCG |
| Slc7a1 | Slc7a1_sg2 | CACAAACGTGAAATACGGTG |
| Slc7a1 | Slc7a1_sg3 | TGACGTGAGAACTCTCCGAT |
| Slc7a1 | Slc7a1_sg4 | CCAGGTCCTTCAGTTCAAAG |
| Smox | Smox_sg1 | CGAGAGTCAGAACAGCGTCG |
| Smox | Smox_sg2 | CAACTCGCATGAAGCCCGAG |
| Smox | Smox_sg3 | GCTCGATCTCAGGACCCCGG |
| Smox | Smox_sg4 | GCCTGCTACCTTACCAACCG |
| Sod1 | Sod1_sg1 | CAGTATGGGGACAATACACA |
| Sod1 | Sod1_sg2 | GACTGCTGGAAAGGACGGTG |
| Sod1 | Sod1_sg3 | TAAGAAACATGGTGGCCCGG |
| Sod1 | Sod1_sg4 | AAAGCGGTGTGCGTGCTGAA |
| Sod2 | Sod2_sg1 | GGCGTTGAGATTGTTCACGT |
| Sod2 | Sod2_sg2 | ATGATCTGCGCGTTAATGTG |
| Sod2 | Sod2_sg3 | ACAAACCTGAGCCCTAAGGG |
| Sod2 | Sod2_sg4 | CCTGCACTGAAGTTCAATGG |
| Sptlc1 | Sptlc1_sg1 | AATGTGCCATAGAACCCTCG |
| Sptlc1 | Sptlc1_sg2 | CCCTCCAACCCACAACATCG |
| Sptlc1 | Sptlc1_sg3 | TCCTGCGTACTCTAAGAGAG |
| Sptlc1 | Sptlc1_sg4 | TTTGTCGTAGAATCCTCGCA |
| Sptlc2 | Sptlc2_sg1 | GTTGTGTTTGAAGATTCGAA |
| Sptlc2 | Sptlc2_sg2 | TGAGAGCAATCACTTCAGGA |
| Sptlc2 | Sptlc2_sg3 | AATCTCGAAGATATCCAAAG |
| Sptlc2 | Sptlc2_sg4 | ACAACTATCTTGGATTTGCG |
| Sptssa | Sptssa_sg1 | CAGGTACTGGTAGTAGAACC |
| Sptssa | Sptssa_sg2 | CTGAACACGGTTCGCTCCCA |
| Sptssa | Sptssa_sg3 | CCATCACGCAGATTCGATGC |
| Sptssa | Sptssa_sg4 | GTACAGGGCCATCCCCACCA |
| Tbx21 | Tbx21_sg1 | AGTCTGGGTGGACATATAAG |
| Tbx21 | Tbx21_sg2 | AGGACTACGCATTGCCCGCG |
| Tbx21 | Tbx21_sg3 | GACCCGACCGATCGCCGCGC |
| Tbx21 | Tbx21_sg4 | GGCTTCCAACAATGTGACCC |
| Uqcrh | Uqcrh_sg1 | TGGATCTGGAGACCCCAAAG |
| Uqcrh | Uqcrh_sg2 | GACGAACGAAAGATGCTCAC |
| Uqcrh | Uqcrh_sg3 | AATCCTCTTCTGTCTGTGAC |
| Uqcrh | Uqcrh_sg4 | GCTCTCTCACTGTTGTTAGG |

| | | |
|---|---|---|
| Tsc2 | Tsc2_sg1 | TGAACCACATGGCTATGACG |
| Tsc2 | Tsc2_sg2 | CACAGGGTGATAATGAACAG |
| Tsc2 | Tsc2_sg3 | CAGCTCCAAAGACCCTTGAG |
| Tsc2 | Tsc2_sg4 | CTGATCCTAGCACACATGTG |
| Aldoa | Aldoa_sg1 | AATGGCGAGACAACTACCCA |
| Aldoa | Aldoa_sg2 | CCTTGCCCGGAGCCACAATG |
| Aldoa | Aldoa_sg3 | CCACGAGACACTGTACCAGA |
| Aldoa | Aldoa_sg4 | GCCAGCATCTGCCAGCAGGT |
| NTC | NTC_sg1 | AAAAAGTCCGCGATTACGTC |
| NTC | NTC_sg2 | AAAACGGCTCGATCGGTGAT |
| NTC | NTC_sg3 | AAAACGTAATTATACCGAGC |
| NTC | NTC_sg4 | AAAATTGCACCTTCCCGGCC |
| NTC | NTC_sg5 | AAACCCCCGCGCGGAGCGTC |
| NTC | NTC_sg6 | AAACCTAGCGTAGATTCGGC |
| NTC | NTC_sg7 | AAACGAGGCTGTTCGTACAC |
| NTC | NTC_sg8 | AAACTCATACGTAGCGAATC |

### Supplementary Table 3. NGS PCR2 indexing primers for pooled CRISPR screen sequencing

| Primer Name | Full Primer Sequence (5'→3') | Adapter Type | Index Barcode Sequence | Notes |
|---|---|---|---|---|
| LenNGS-Fwd-1 | AATGATACGGCGACCACCGAGATCTACACTCTTTCCCTACACGACGCTCTTCCGATCTTAAGTAGAGGCTTTATATATCTTGTGGAAAGGACGAAACACC | P5 | TAAGTAGAG | Forward indexing primer |
| LenNGS-Fwd-2 | AATGATACGGCGACCACCGAGATCTACACTCTTTCCCTACACGACGCTCTTCCGATCTATCATGCTTAGCTTTATATATCTTGTGGAAAGGACGAAACACC | P5 | ATCATGCTTA | Forward indexing primer |
| LenNGS-Fwd-3 | AATGATACGGCGACCACCGAGATCTACACTCTTTCCCTACACGACGCTCTTCCGATCTGATGCACATCTGCTTTATATATCTTGTGGAAAGGACGAAACACC | P5 | GATGCACATCT | Forward indexing primer |
| LenNGS-Fwd-4 | AATGATACGGCGACCACCGAGATCTACACTCTTTCCCTACACGACGCTCTTCCGATCTCGATTGCTCGACGCTTTATATATCTTGTGGAAAGGACGAAACACC | P5 | CGATTGCTCGAC | Forward indexing primer |
| LenNGS-Fwd-5 | AATGATACGGCGACCACCGAGATCTACACTCTTTCCCTACACGACGCTCTTCCGATCTTCGATAGCAATTCGCTTTATATATCTTGTGGAAAGGACGAAACACC | P5 | TCGATAGCAATTC | Forward indexing primer |
| LenNGS-Fwd-6 | AATGATACGGCGACCACCGAGATCTACACTCTTTCCCTACACGACGCTCTTCCGATCTATCGATAGTTGCTTGCTTTATATATCTTGTGGAAAGGACGAAACACC | P5 | ATCGATAGTTGCTT | Forward indexing primer |
| LenNGS-Fwd-7 | AATGATACGGCGACCACCGAGATCTACACTCTTTCCCTACACGACGCTCTTCCGATCTGATCGATCCAGTTAGGCTTTATATATCTTGTGGAAAGGACGAAACACC | P5 | GATCGATCCAGTTAG | Forward indexing primer |
| LenNGS-Fwd-8 | AATGATACGGCGACCACCGAGATCTACACTCTTTCCCTACACGACGCTCTTCCGATCTCGATCGATTTGAGCCTGCTTTATATATCTTGTGGAAAGGACGAAACACC | P5 | CGATCGATTTGAGCCT | Forward indexing primer |
| LenNGS-Fwd-9 | AATGATACGGCGACCACCGAGATCTACACTCTTTCCCTACACGACGCTCTTCCGATCTACGATCGATACACGATCGCTTTATATATCTTGTGGAAAGGACGAAACACC | P5 | ACGATCGATACACGATC | Forward indexing primer |
| LenNGS-Fwd-10 | AATGATACGGCGACCACCGAGATCTACACTCTTTCCCTACACGACGCTCTTCCGATCTTACGATCGATGGTCCAGAGCTTTATATATCTTGTGGAAAGGACGAAACACC | P5 | TACGATCGATGGTCCAGA | Forward indexing primer |
| LenNGS-Rev-1 | CAAGCAGAAGACGGCATACGAGATAAGTAGAGGTGACTGGAGTTCAGACGTGTGCTCTTCCGATCTCCGACTCGGTGCCACTTTTTCAA | P7 | AAGTAGAG | Reverse indexing primer |
| LenNGS-Rev-2 | CAAGCAGAAGACGGCATACGAGATACACGATCGTGACTGGAGTTCAGACGTGTGCTCTTCCGATCTCCGACTCGGTGCCACTTTTTCAA | P7 | ACACGATC | Reverse indexing primer |
| LenNGS-Rev-3 | CAAGCAGAAGACGGCATACGAGATCGCGCGGTGTGACTGGAGTTCAGACGTGTGCTCTTCCGATCTCCGACTCGGTGCCACTTTTTCAA | P7 | CGCGCGGT | Reverse indexing primer |
| LenNGS-Rev-4 | CAAGCAGAAGACGGCATACGAGATCATGATCGGTGACTGGAGTTCAGACGTGTGCTCTTCCGATCTCCGACTCGGTGCCACTTTTTCAA | P7 | CATGATCG | Reverse indexing primer |
| LenNGS-Rev-5 | CAAGCAGAAGACGGCATACGAGATCGTTACCAGTGACTGGAGTTCAGACGTGTGCTCTTCCGATCTCCGACTCGGTGCCACTTTTTCAA | P7 | CGTTACCA | Reverse indexing primer |
| LenNGS-Rev-6 | CAAGCAGAAGACGGCATACGAGATTCCTTGGTGTGACTGGAGTTCAGACGTGTGCTCTTCCGATCTCCGACTCGGTGCCACTTTTTCAA | P7 | TCCTTGGT | Reverse indexing primer |
| LenNGS-Rev-7 | CAAGCAGAAGACGGCATACGAGATAACGCATTGTGACTGGAGTTCAGACGTGTGCTCTTCCGATCTCCGACTCGGTGCCACTTTTTCAA | P7 | AACGCATT | Reverse indexing primer |
| LenNGS-Rev-8 | CAAGCAGAAGACGGCATACGAGATACAGGTATGTGACTGGAGTTCAGACGTGTGCTCTTCCGATCTCCGACTCGGTGCCACTTTTTCAA | P7 | ACAGGTAT | Reverse indexing primer |
| LenNGS-Rev-9 | CAAGCAGAAGACGGCATACGAGATAGGTAAGGGTGACTGGAGTTCAGACGTGTGCTCTTCCGATCTCCGACTCGGTGCCACTTTTTCAA | P7 | AGGTAAGG | Reverse indexing primer |
| LenNGS-Rev-10 | CAAGCAGAAGACGGCATACGAGATAACAATGGGTGACTGGAGTTCAGACGTGTGCTCTTCCGATCTCCGACTCGGTGCCACTTTTTCAA | P7 | AACAATGG | Reverse indexing primer |
| LenNGS-Rev-11 | CAAGCAGAAGACGGCATACGAGATACTGTATCGTGACTGGAGTTCAGACGTGTGCTCTTCCGATCTCCGACTCGGTGCCACTTTTTCAA | P7 | ACTGTATC | Reverse indexing primer |
| LenNGS-Rev-12 | CAAGCAGAAGACGGCATACGAGATAGGTCGCAGTGACTGGAGTTCAGACGTGTGCTCTTCCGATCTCCGACTCGGTGCCACTTTTTCAA | P7 | AGGTCGCA | Reverse indexing primer |
| LenNGS-Rev-13 | CAAGCAGAAGACGGCATACGAGATTCTCATGAGTGACTGGAGTTCAGACGTGTGCTCTTCCGATCTCCGACTCGGTGCCACTTTTTCAA | P7 | TCTCATGA | Reverse indexing primer |
| LenNGS-Rev-14 | CAAGCAGAAGACGGCATACGAGATCTCTGCAGGTGACTGGAGTTCAGACGTGTGCTCTTCCGATCTCCGACTCGGTGCCACTTTTTCAA | P7 | CTCTGCAG | Reverse indexing primer |
| LenNGS-Rev-15 | CAAGCAGAAGACGGCATACGAGATCACTATCAGTGACTGGAGTTCAGACGTGTGCTCTTCCGATCTCCGACTCGGTGCCACTTTTTCAA | P7 | CACTATCA | Reverse indexing primer |
| LenNGS-Rev-16 | CAAGCAGAAGACGGCATACGAGATTGTCGCTGGTGACTGGAGTTCAGACGTGTGCTCTTCCGATCTCCGACTCGGTGCCACTTTTTCAA | P7 | TGTCGCTG | Reverse indexing primer |
| LenNGS-Rev-17 | CAAGCAGAAGACGGCATACGAGATGCAGAGCTGTGACTGGAGTTCAGACGTGTGCTCTTCCGATCTCCGACTCGGTGCCACTTTTTCAA | P7 | GCAGAGCT | Reverse indexing primer |
| LenNGS-Rev-18 | CAAGCAGAAGACGGCATACGAGATATGAGATCGTGACTGGAGTTCAGACGTGTGCTCTTCCGATCTCCGACTCGGTGCCACTTTTTCAA | P7 | ATGAGATC | Reverse indexing primer |
| LenNGS-Rev-19 | CAAGCAGAAGACGGCATACGAGATTTGCTGCCGTGACTGGAGTTCAGACGTGTGCTCTTCCGATCTCCGACTCGGTGCCACTTTTTCAA | P7 | TTGCTGCC | Reverse indexing primer |
| LenNGS-Rev-20 | CAAGCAGAAGACGGCATACGAGATCCATCATTGTGACTGGAGTTCAGACGTGTGCTCTTCCGATCTCCGACTCGGTGCCACTTTTTCAA | P7 | CCATCATT | Reverse indexing primer |

**Supplementary Table 4. RT-PCR primers used in study.**

| Gene | Primer Name | Direction | Sequence (5'→3') |
|---|---|---|---|
| Cpt1a | Mus_Cpt1a_Fwd | Forward | AGTGGCCTCACAGACTCCAG |
| Cpt1a | Mus_Cpt1a_Rev | Reverse | GCCCATGTTGTACAGCTTCC |
| Rpl37 | Mus_Rpl37_Fwd | Forward | CTACCGCAGATTCAGACATGGA |
| Rpl37 | Mus_Rpl37_Rev | Reverse | ACCGAACTCTGAACCGATGT |
| Gapdh | Mus_Gapdh_Fwd | Forward | TCAGGAGAGTGTTTCCTCGTC |
| Gapdh | Mus_Gapdh_Rev | Reverse | TTTGCCGTGAGTGGAGTCAT |
| Lcn2 | Mus_Lcn2_Fwd | Forward | ACTCTGGGAAATATGCACAGGTAT |
| Lcn2 | Mus_Lcn2_Rev | Reverse | AAGCGGGTGAAACGTTCCTT |
| Slpi | Mus_Slpi_Fwd | Forward | TTGAGAAGCCACAATGCCGT |
| Slpi | Mus_Slpi_Rev | Reverse | GAGTTTTGACGCACCTCCCA |
| Cxcl5 | Mus_Cxcl5_Fwd | Forward | CCCTACGGTGGAAGTCATAGC |
| Cxcl5 | Mus_Cxcl5_Rev | Reverse | TTAGCTTTCTTTTTGTCACTGCCC |
| Rsad2 | Mus_Rsad2_Fwd | Forward | GTGCCTGAATCTAACCAGAAGATGA |
| Rsad2 | Mus_Rsad2_Rev | Reverse | ATACTTTCCGCCACGCTTCA |
| Cmpk2 | Mus_Cmpk2_Fwd | Forward | GGTAAGACCACACTGACGCA |
| Cmpk2 | Mus_Cmpk2_Rev | Reverse | AGCCACGAGATAATTGCCCA |
| Ifi44 | Mus_Ifi44_Fwd | Forward | GGTACAGACTCTTCACACAGACTT |
| Ifi44 | Mus_Ifi44_Rev | Reverse | TTCTGCACACTCGCCTTGTA |
| Irf7 | Mus_Irf7_Fwd | Forward | CAAGAGAAAATGCTGGGCTCC |
| Irf7 | Mus_Irf7_Rev | Reverse | ATAGGGTTCCTCGTAAACACGG |
| Tmem173 | Mus_Tmem173_Fwd | Forward | GCTGCTGATGCCATACTCCAA |
| Tmem173 | Mus_Tmem173_Rev | Reverse | AGTAGTCCAAGTTCGTGCGAG |
| Ppif | Mus_Ppif_Fwd | Forward | GATGTCGTGCCAAAGACTGC |
| Ppif | Mus_Ppif_Rev | Reverse | AGTGTGAAGTTCTCGTCGGG |
| mtD-loop1 | Mus_mtD-loop1_Fwd | Forward | AATCTACCATCCTCCGTGAAACC |
| mtD-loop1 | Mus_mtD-loop1_Rev | Reverse | TCAGTTTAGCTACCCCCAAGTTTAA |
| mtD-loop2 | Mus_mtD-loop2_Fwd | Forward | CCCTTCCCCATTTGGTCT |
| mtD-loop2 | Mus_mtD-loop2_Rev | Reverse | TGGTTTCACGGAGGATGG |
| mtD-loop3 | Mus_mtD-loop3_Fwd | Forward | TCCTCCGTGAAACCAACAA |
| mtD-loop3 | Mus_mtD-loop3_Rev | Reverse | AGCGAGAAGAGGGGCATT |
| mt16S | Mus_mt16S_Fwd | Forward | CACTGCCTGCCCAGTGA |
| mt16S | Mus_mt16S_Rev | Reverse | ATACCGCGGCCGTTAAA |
| mtNd4 | Mus_mtNd4_Fwd | Forward | AACGGATCCACAGCCGTA |
| mtNd4 | Mus_mtNd4_Rev | Reverse | AGTCCTCGGGCCATGATT |

**Supplementary Table 5. Flow cytometry antibodies used in study.**

| Antibody | Clone | Dilution | Company | Catalog # |
|---|---|---|---|---|
| APC/Cy7-CD45 | 30-F11 | 1:500 | BD Biosciences | 557659 |
| V450-CD45.2 | 104 | 1:500 | Tonbo/Cytek | 75-0454 |
| PE/Cy7-TCRb | H57-597 | 1:200 | Tonbo/Cytek | 60-5961 |
| PerCP/Cy5.5-TCRb | H57-597 | 1:250 | Tonbo/Cytek | 65-5961 |
| RedFluor710-CD8 | 53-6.7 | 1:500 | Tonbo/Cytek | 80-0081 |
| BV605-CD8 | 53-6.7 | 1:400 | Biolegend | 100743 |
| PE/Dazzle594-CD4 | GK1.5 | 1:500 | Biolegend | 100455 |
| PerCP/Cy5.5-CD11b | M1/70 | 1:500 | Tonbo/Cytek | 65-0112 |
| APC/Cy7-CD25 | PC61 | 1:200 | Biolegend | 102025 |
| PerCP/Cy5.5-CD127 | A7R34 | 1:100 | eBioscience | 45-1271 |
| V450-IFNg | XMG1.2 | 1:100 | Tonbo/Cytek | 75-7311 |
| APC-TNFa | MP6-XT22 | 1:50 | eBioscience | 17-7321-82 |
| PE-GATA3 | TWAJ | 1:100 | eBioscience | Dec-66 |
| eFluor660-FoxP3 | FJK-16s | 1:100 | eBioscience | 50-5773-82 |
| APC-IgG1,k | eBRG1 | 1:50 | eBioscience | 17-4301-82 |
